# Cell-specific effects of acute sleep deprivation on transcriptomic signatures of non-neuronal cells in the mouse hippocampus and prefrontal cortex

**DOI:** 10.64898/2026.01.29.702391

**Authors:** Junko Kasuya, Yutong Wang, Yann Vanrobaeys, Marcos Frank, Ted Abel

**Affiliations:** Department of Neuroscience and Pharmacology; Iowa Neuroscience Institute; Graduate Program in Pharmacology; Interdisciplinary Graduate Program in Genetics, Carver College of Medicine, University of Iowa, Iowa, USA; Department of Translational Medicine and Physiology, Elson S. Floyd College of Medicine, Washington State University, Spokane, WA, USA; Gleason Institute for Neuroscience, Washington State University, Spokane, WA, USA; Sleep Performance and Research Center, Elson S. Floyd College of Medicine, Washington State University, Spokane, WA, USA; Bioinformatics & Computing Core Facility of the Swanson Biotechnology Center, Koch Institute for Integrative Cancer Research, Massachusetts Institute of Technology, 77 Massachusetts Ave. 68-304d, Cambridge, MA, USA

**Author notes:** Corresponding author ADDRESS FOR CORRESPONDENCE: Ted Abel, Ph.D. Iowa Neuroscience Institute, Department of Neuroscience and Pharmacology, Carver College of Medicine, University of Iowa, 2312 Pappajohn Biomedical Discovery Building, 169 Newton Road, Iowa City, Iowa 52242-1903, United States.

**Keywords:** acute sleep deprivation, hippocampus, prefrontal cortex, non-neuronal cells, single nuclei RNAseq, astrocytes, cholesterol synthesis, mitochondria, cadherin 2, primary cilia, oligodendrocytes

## Abstract

Sleep is essential for maintaining cognitive, emotional, metabolic, and immune functions. Although research on sleep homeostasis is traditionally neuron-centric, increasing evidence indicates non-neuronal cells also play critical roles. In this study, we performed transcriptomic analyses of non-neuronal nuclei from the hippocampus and prefrontal cortex of mice subjected to acute sleep deprivation (SD). We found that acute SD induces robust, cell-type- and region-specific transcriptional reprogramming in astrocytes and oligodendrocytes. The most pronounced changes occurred in astrocytes, including downregulation of cholesterol biosynthesis genes in both brain regions, accompanied by region-specific and opposing regulation of genes involved in mitochondrial function and neurodegeneration-related pathways. Notably, genes associated with primary cilia were selectively induced in cortical astrocytes. In oligodendrocytes, acute SD led to downregulation of genes encoding cell-adhesion molecules. Together, these findings provide molecular evidence that non-neuronal cells actively contribute to sleep regulation and suggest novel potential mechanisms that broaden our understanding of sleep homeostasis.

## Introduction

Sleep is a well-regulated and reversible state defined by distinct behavioral, homeostatic, and electrophysiological properties. Sleep plays a critical role in various biological functions, including cognitive processes, immune system performance, physiological and neural homeostasis, and even survival. Sleep deprivation, a growing concern in modern society, has been linked to numerous health issues^1,2^. In humans, one of the most notable consequences of sleep deprivation is a significant decline in attention and memory. Research shows that insufficient sleep—whether acute or chronic—can impair performance on tasks requiring sustained attention or prolonged focus on repetitive stimuli^3,4^. Sleep disturbances, often categorized as insomnia, are also closely associated with a wide range of medical conditions, either as a contributing factor or a consequence. These conditions include obesity, diabetes, metabolic and neuroendocrine disorders, as well as psychiatric and neurological illnesses^5,6^. Despite the prevalence and impact of sleep disorders, current treatments are limited. This underscores the importance of understanding the mechanisms underlying sleep regulation and sleep function, which is essential for developing more effective therapies.

Research into the molecular and cellular mechanisms underlying sleep homeostasis has traditionally focused on neurons, largely because different sleep stages were initially characterized using EEG, a technique that reflects neuronal activity. However, the role of non-neuronal cells in sleep regulation and function was proposed as early as the work of Santiago Ramón y Cajal over a century ago^7^. In recent years, this hypothesis has been supported by accumulating evidence^8–11^. Although astrocytes are not electrically excitable, they influence neuronal activity through mechanisms such as glutamate uptake and release, maintenance of energy metabolism, phagocytosis, and ATP release, which serves as a source of extracellular adenosine^11–16^. A recent study in fruit flies demonstrated that glial cells, including astrocytes, play a critical role during sleep by clearing metabolites generated by neurons^16^. This finding further supports the concept of neuron–astrocyte neurometabolic coupling (NMC) as essential for sleep regulation^15^. Intracellular Ca²⁺ levels in astrocytes correlate with changes in sleep pressure, and genetically reducing glial Ca²⁺ levels impairs the homeostatic response to sleep deprivation^17^. Furthermore, activating Gq-protein signaling in astrocytes in the basal forebrain prolongs wakefulness without increasing sleep need^18^. Another major glial cell type in the mature adult brain is oligodendrocytes. These cells produce myelin, an electrical insulator essential for proper neural activity^19^. Recent studies suggest that experience-dependent myelin plasticity in the adult brain also plays a crucial role in maintaining normal nervous system function, acting alongside neuronal plasticity^20,21^. Consistent with this idea, gene expression studies examining the cellular responses of oligodendrocytes to sleep and sleep loss have identified genes encoding myelin structural proteins and associated enzymes^22^.

Sleep homeostasis and the functional role of sleep are regulated through the coordinated actions of various cell types across multiple brain regions, mediated by distinct molecular pathways. Like other complex biological phenomena, these processes involve intricate interactions between neurons and non-neuronal cells. In the current study, we investigated the molecular effects of acute sleep deprivation on non-neuronal cells—specifically astrocytes and mature oligodendrocytes—using a single-nuclear RNA sequencing approach. We examined the hippocampus (HPC) and prefrontal cortex (PFC), two brain regions critically affected by sleep deprivation. The hippocampus is essential for memory and learning, which are impaired by sleep loss^23–26^, while the PFC is closely linked to sleep homeostasis^18,27^, as well as higher cognitive functions^28^.

We report the biological pathways affected by acute SD at cell type level, which are identified based on the molecular changes from the snRNAseq analyses. Those include genes encoding proteins involved in cholesterol biosynthesis which were downregulated in astrocytes in both the HPC and PFC, with a more pronounced effect in the PFC. In addition, biological pathways associated with neurodegeneration, in particular, the mitochondrial electron transport chain, were altered in astrocytes across both regions, but in opposite directions. The transcriptomic changes in cortical astrocytes also include genes linked to the assembly and elongation of primary cilia. In mature oligodendrocytes (MOLs), we observed a significant and MOL-specific downregulation of cell adhesion molecules including Cadherin 2 (Cdh2), suggesting a synergistic effect with reduced cholesterol synthesis in compromising synaptic stability.

## RESULTS

### Hippocampus and prefrontal cortex present similar cell type profiles

Single-cell or single-nucleus RNA sequencing (RNAseq) has been instrumental in uncovering the detailed transcriptomic signatures of specific cell types. To overcome the limitations in analysis depth, often constrained by the number of nuclei that can be processed at a time, we enriched a pool of nuclei for non-neuronal types to be used as the starting material. Using the *Aldh1l1*-cre, we labeled glial cells with a Sun1/sfGFP^29^ nuclear-tag. However, more than half of the tagged nuclei also expressed *NeuN* (*Rbfox3*), a general neuronal marker. To refine our approach, we excluded the subpopulation of tagged nuclei that expressed *NeuN* using fluorescence-activated nuclei sorting (FANS) (Figure S1A-D). We designated the remaining pool of nuclei—those nuclei expressing *Aldh1l1*-cre-driving Sun1/sfGFP but lacking *NeuN* expression—as the “non-neuronal” nuclei population, which was subsequently used as the starting material for snRNAseq analyses. This population comprised 31.8% and 23.2% of the total nuclei in the hippocampus (HPC) and prefrontal cortex (PFC), respectively (Figure S1C and S1D). This strategy enabled us to significantly increase the recovery of nuclei from non-neuronal cell types, such as astrocytes, mature oligodendrocytes (MOL), endothelial cells, and oligodendrocyte progenitor cells (OPCs) (Figure S2A-C).

The non-neuronal nuclear pool was prepared from the HPC or PFC of acutely sleep deprived (SD), or home-caged non sleep deprived (NSD) animals as described in the method section. UMAP of nuclei from both brain regions showed similar cell profiles, with astrocytes and MOLs being the two major components (Figure 1A and 1B). In the UMAP, both astrocytes and MOLs were seen subdivided into two subtypes, labeled as -1 and -2, both of which expressed typical cell markers for astrocytes and MOLs, respectively, but showed distinct transcriptional signatures (Supplemental Data). The molecular responses to acute SD in both subtypes were, however, largely similar, so they were combined for further analysis and are referred to as astrocytes or MOLs hereafter. In addition to the two major astrocyte subtypes, a small additional population of astrocytes was identified in both HPC and PFC. This subset of astrocytes expresses genes encoding the SNARE proteins which perform exocytosis, such as Snap25, Syt7, and Vamp2, in addition to astrocyte marker genes, and also are also enriched in glutamate signaling and Ca^2+^ signaling pathways (Figure S3A and S3B, Supplemental Data), functions that resembles the mechanism of gliotransmission^30–32^.

**Figure 1.**
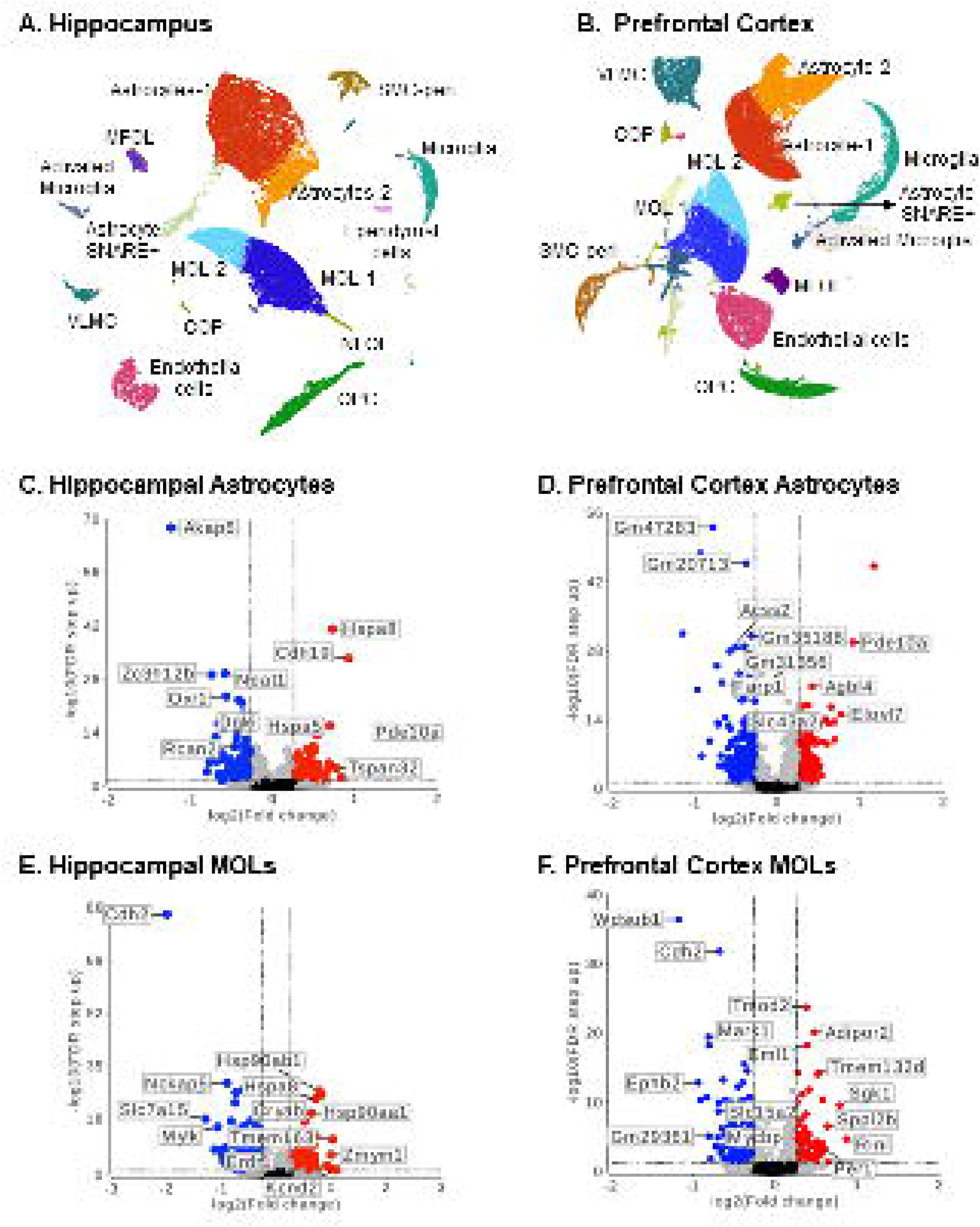
Single-nuclear transcriptome analyses of non-neuronal cells from mice hippocampus and prefrontal cortex after acute SD. (**A** and **B**) Dimension reduction UMAP of non-neuronal nuclei from mice hippocampus **(A)** or prefrontal cortex **(B)**. (**C** - **F**) Volcano plot of DEGs identified in response to acute sleep deprivation in astrocytes **(C and D)** or in mature oligodendrocytes (MOLs) **(E and F)** from HPC **(C and E)** or PFC **(D and F)**. The red color indicates upregulated DEGs, the blue color represents downregulated DEGs, and the gray color means genes with no significance. The criteria the significance is | FC | >1.2 and adjusted p value <0.05.

### Astrocytes in HPC and PFC respond differently to acute SD

Differentially expressed genes (DEGs) were identified using a fold-change cutoff of >1.2 or <-1.2 and an FDR < 0.05, yielding 532 and 486 DEGs in HPC and PFC, respectively. DEG lists for each region are provided in the Supplemental Data and visualized as volcano plots (Figure 1C and 1E for HPC and Figure 1D and 1F for PFC, representing astrocytes and MOL, respectively). 74 astrocytic DEGs overlapped between the two regions (Figure S4A and S4B).

These DEGs were used for identifying biological pathways affected by acute SD using the Database for Annotation, Visualization, and Integrated Discovery (DAVID). Acute SD significantly enriched 37 and 51 biological pathway or molecular function terms in HPC and PFC astrocytes, respectively (Benjamini-adjusted p < 0.05). To simplify interpretation, these terms were clustered by functional similarity (Table 1) and represented as fractional distributions in Figure 2. Pathways not clustered were grouped as “others” and listed in the Table 1.

**Table 1.**
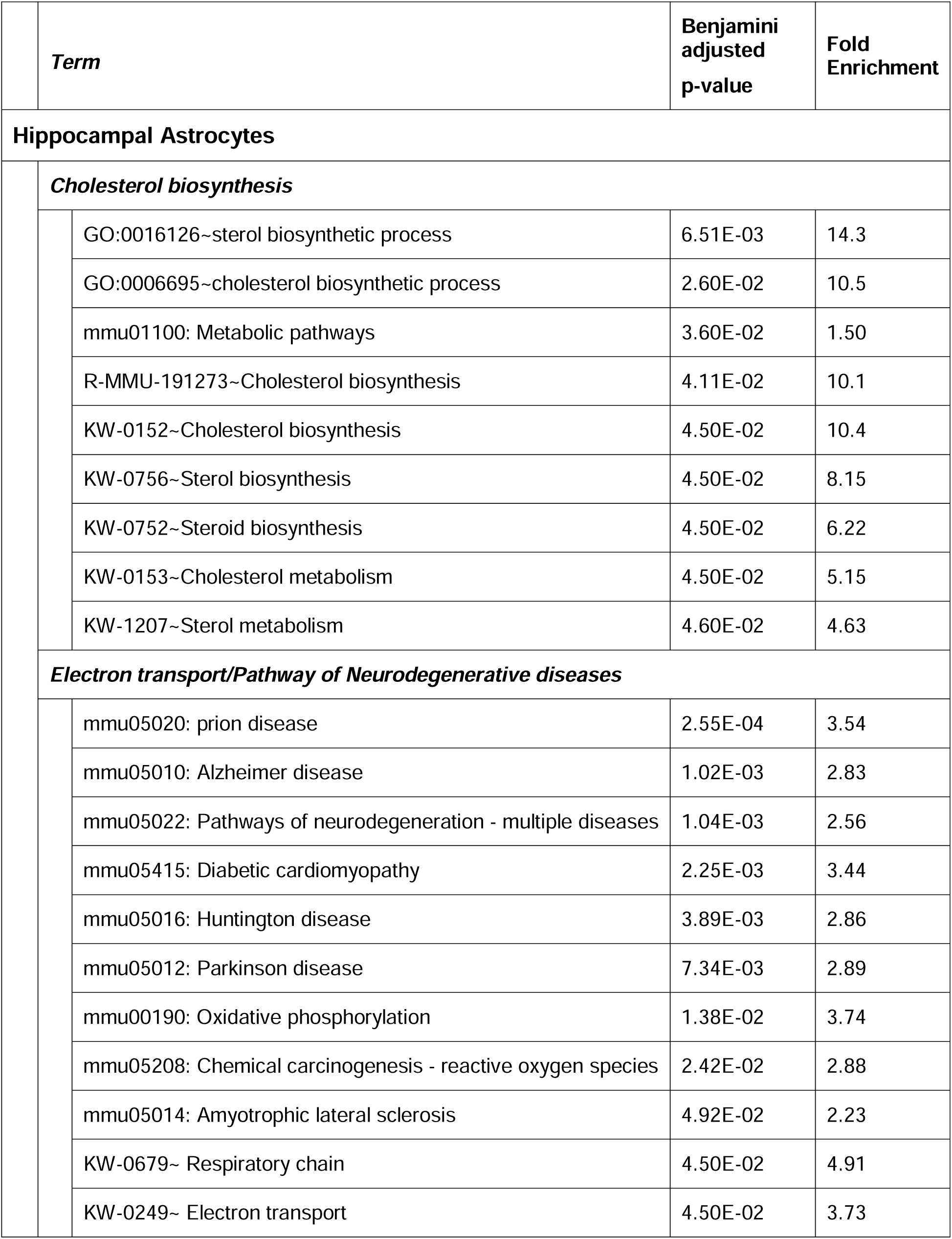
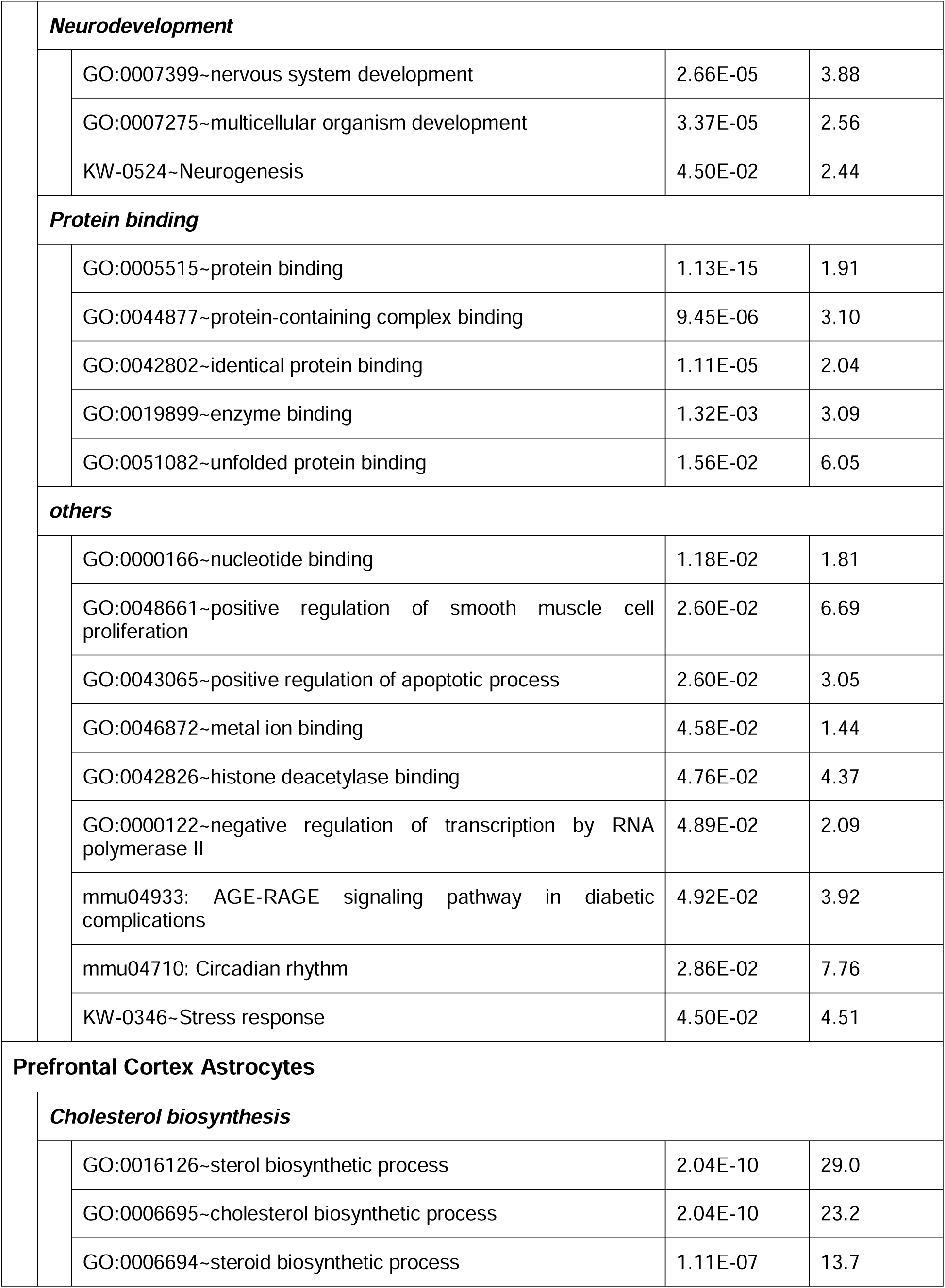
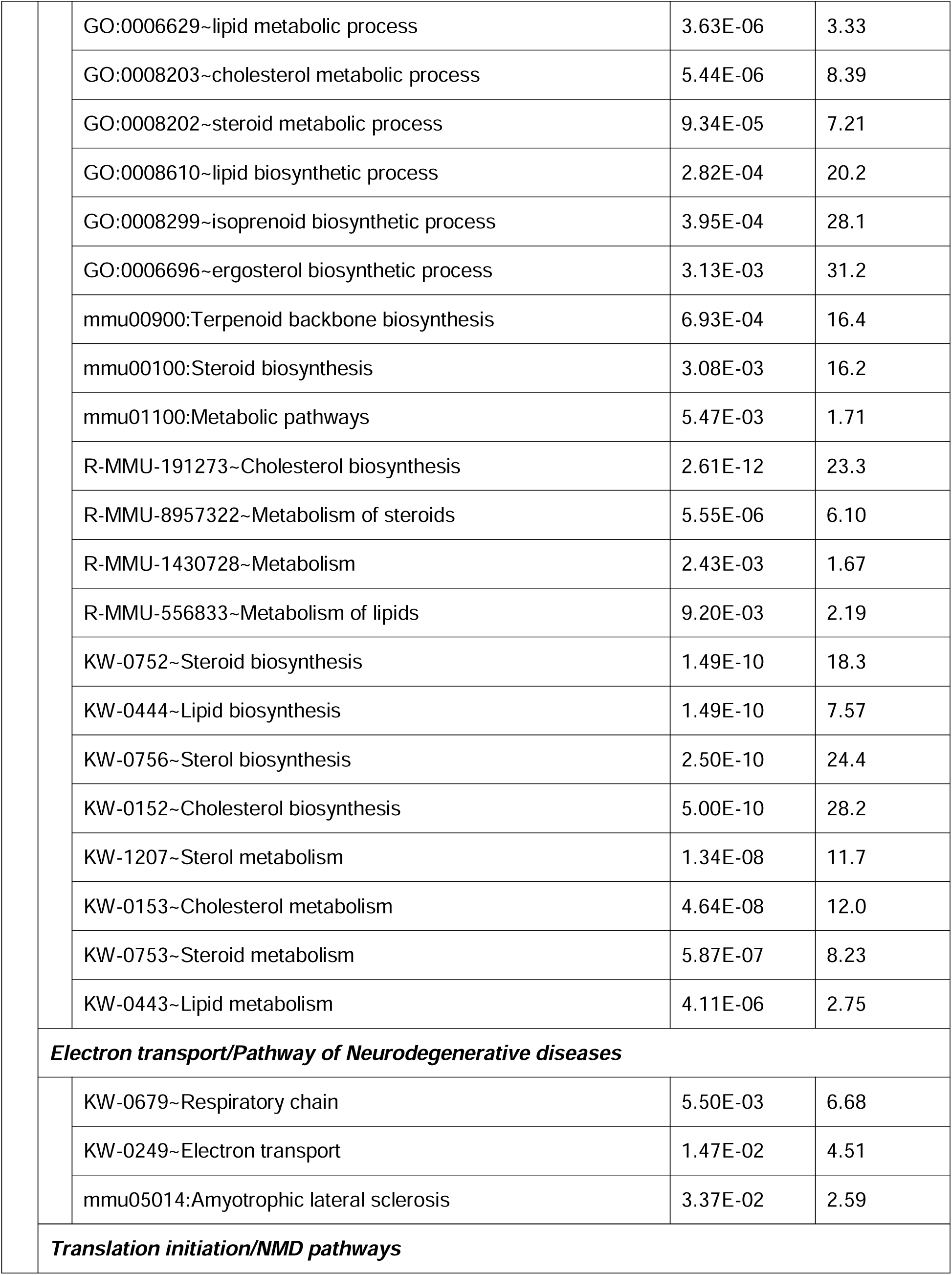
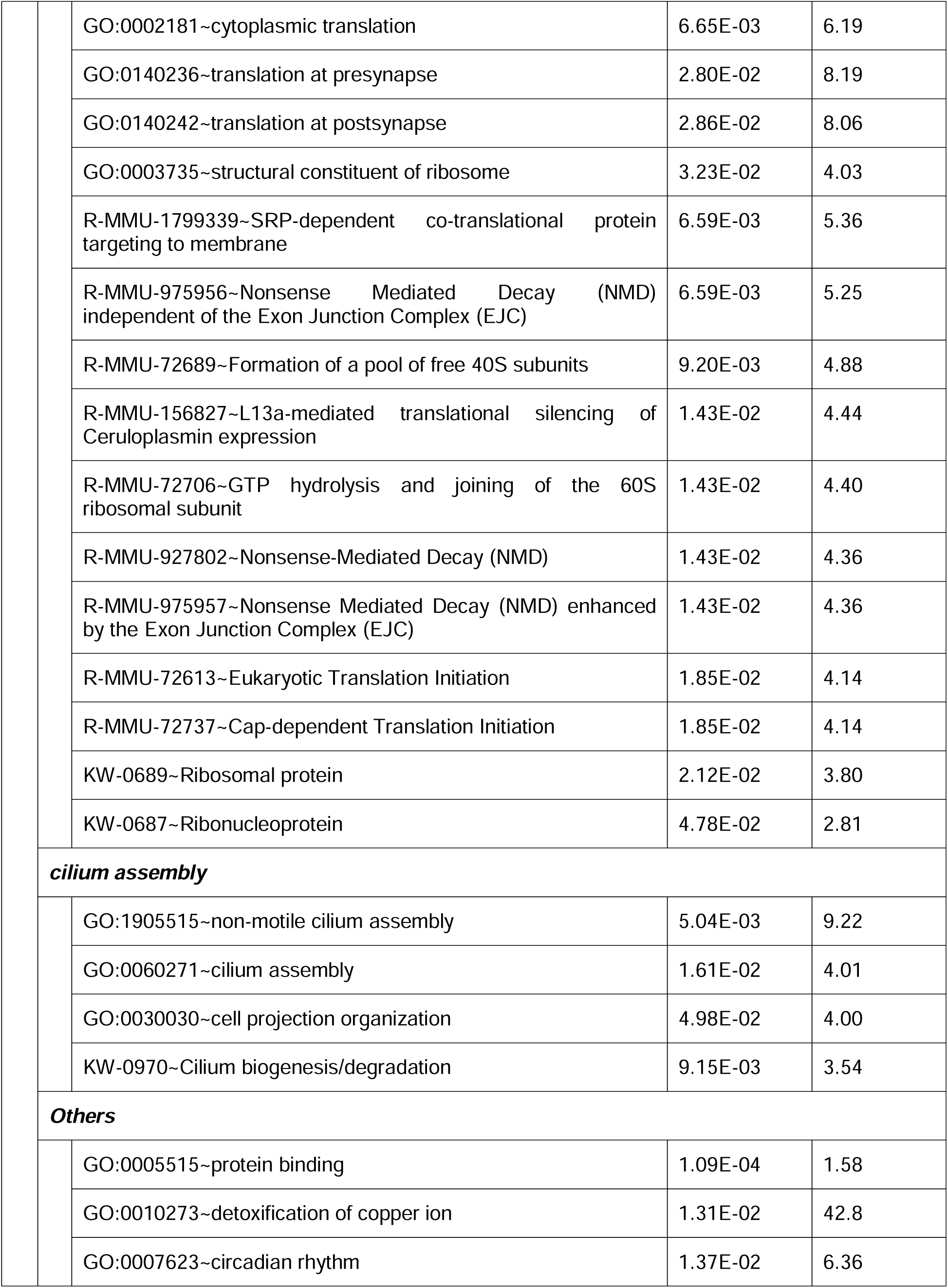
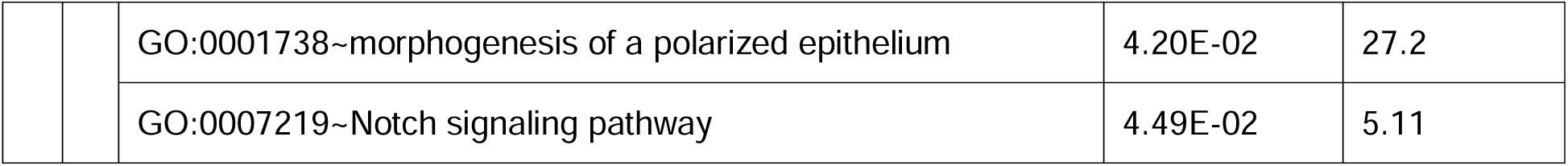
Enhanced biological pathways in Astrocytes

|  | *Term* | **Benjamini**<br>**adjusted**<br>**p-value** | **Fold**<br>**Enrichment** |
| --- | --- | --- | --- |
| **Hippocampal Astrocytes** |  |  |  |
|  | ***Cholesterol biosynthesis*** |  |  |
|  | GO:0016126~sterol biosynthetic process | 6.51E-03 | 14.3 |
|  | GO:0006695~cholesterol biosynthetic process | 2.60E-02 | 10.5 |
|  | mmu01100: Metabolic pathways | 3.60E-02 | 1.50 |
|  | R-MMU-191273~Cholesterol biosynthesis | 4.11E-02 | 10.1 |
|  | KW-0152~Cholesterol biosynthesis | 4.50E-02 | 10.4 |
|  | KW-0756~Sterol biosynthesis | 4.50E-02 | 8.15 |
|  | KW-0752~Steroid biosynthesis | 4.50E-02 | 6.22 |
|  | KW-0153~Cholesterol metabolism | 4.50E-02 | 5.15 |
|  | KW-1207~Sterol metabolism | 4.60E-02 | 4.63 |
|  | ***Electron transport/Pathway of Neurodegenerative diseases*** |  |  |
|  | mmu05020: prion disease | 2.55E-04 | 3.54 |
|  | mmu05010: Alzheimer disease | 1.02E-03 | 2.83 |
|  | mmu05022: Pathways of neurodegeneration - multiple diseases | 1.04E-03 | 2.56 |
|  | mmu05415: Diabetic cardiomyopathy | 2.25E-03 | 3.44 |
|  | mmu05016: Huntington disease | 3.89E-03 | 2.86 |
|  | mmu05012: Parkinson disease | 7.34E-03 | 2.89 |
|  | mmu00190: Oxidative phosphorylation | 1.38E-02 | 3.74 |
|  | mmu05208: Chemical carcinogenesis - reactive oxygen species | 2.42E-02 | 2.88 |
|  | mmu05014: Amyotrophic lateral sclerosis | 4.92E-02 | 2.23 |
|  | KW-0679~ Respiratory chain | 4.50E-02 | 4.91 |
|  | KW-0249~ Electron transport | 4.50E-02 | 3.73 |
| **Neurodevelopment** |  |  |  |
|  | GO:0007399~nervous system development | 2.66E-05 | 3.88 |
|  | GO:0007275~multicellular organism development | 3.37E-05 | 2.56 |
|  | KW-0524~Neurogenesis | 4.50E-02 | 2.44 |
| **Protein binding** |  |  |  |
|  | GO:0005515~protein binding | 1.13E-15 | 1.91 |
|  | GO:0044877~protein-containing complex binding | 9.45E-06 | 3.10 |
|  | GO:0042802~identical protein binding | 1.11E-05 | 2.04 |
|  | GO:0019899~enzyme binding | 1.32E-03 | 3.09 |
|  | GO:0051082~unfolded protein binding | 1.56E-02 | 6.05 |
| **others** |  |  |  |
|  | GO:0000166~nucleotide binding | 1.18E-02 | 1.81 |
|  | GO:0048661~positive regulation of smooth muscle cell proliferation | 2.60E-02 | 6.69 |
|  | GO:0043065~positive regulation of apoptotic process | 2.60E-02 | 3.05 |
|  | GO:0046872~metal ion binding | 4.58E-02 | 1.44 |
|  | GO:0042826~histone deacetylase binding | 4.76E-02 | 4.37 |
|  | GO:0000122~negative regulation of transcription by RNA polymerase II | 4.89E-02 | 2.09 |
|  | mmu04933: AGE-RAGE signaling pathway in diabetic complications | 4.92E-02 | 3.92 |
|  | mmu04710: Circadian rhythm | 2.86E-02 | 7.76 |
|  | KW-0346~Stress response | 4.50E-02 | 4.51 |
| **Prefrontal Cortex Astrocytes** |  |  |  |
| **Cholesterol biosynthesis** |  |  |  |
|  | GO:0016126~sterol biosynthetic process | 2.04E-10 | 29.0 |
|  | GO:0006695~cholesterol biosynthetic process | 2.04E-10 | 23.2 |
|  | GO:0006694~steroid biosynthetic process | 1.11E-07 | 13.7 |
|  | GO:0006629~lipid metabolic process | 3.63E-06 | 3.33 |
|  | GO:0008203~cholesterol metabolic process | 5.44E-06 | 8.39 |
|  | GO:0008202~steroid metabolic process | 9.34E-05 | 7.21 |
|  | GO:0008610~lipid biosynthetic process | 2.82E-04 | 20.2 |
|  | GO:0008299~isoprenoid biosynthetic process | 3.95E-04 | 28.1 |
|  | GO:0006696~ergosterol biosynthetic process | 3.13E-03 | 31.2 |
|  | mmu00900:Terpenoid backbone biosynthesis | 6.93E-04 | 16.4 |
|  | mmu00100:Steroid biosynthesis | 3.08E-03 | 16.2 |
|  | mmu01100:Metabolic pathways | 5.47E-03 | 1.71 |
|  | R-MMU-191273~Cholesterol biosynthesis | 2.61E-12 | 23.3 |
|  | R-MMU-8957322~Metabolism of steroids | 5.55E-06 | 6.10 |
|  | R-MMU-1430728~Metabolism | 2.43E-03 | 1.67 |
|  | R-MMU-556833~Metabolism of lipids | 9.20E-03 | 2.19 |
|  | KW-0752~Steroid biosynthesis | 1.49E-10 | 18.3 |
|  | KW-0444~Lipid biosynthesis | 1.49E-10 | 7.57 |
|  | KW-0756~Sterol biosynthesis | 2.50E-10 | 24.4 |
|  | KW-0152~Cholesterol biosynthesis | 5.00E-10 | 28.2 |
|  | KW-1207~Sterol metabolism | 1.34E-08 | 11.7 |
|  | KW-0153~Cholesterol metabolism | 4.64E-08 | 12.0 |
|  | KW-0753~Steroid metabolism | 5.87E-07 | 8.23 |
|  | KW-0443~Lipid metabolism | 4.11E-06 | 2.75 |
|  | ***Electron transport/Pathway of Neurodegenerative diseases*** |  |  |
|  | KW-0679~Respiratory chain | 5.50E-03 | 6.68 |
|  | KW-0249~Electron transport | 1.47E-02 | 4.51 |
|  | mmu05014:Amyotrophic lateral sclerosis | 3.37E-02 | 2.59 |
|  | ***Translation initiation/NMD pathways*** |  |  |
|  | GO:0002181~cytoplasmic translation | 6.65E-03 | 6.19 |
|  | GO:0140236~translation at presynapse | 2.80E-02 | 8.19 |
|  | GO:0140242~translation at postsynapse | 2.86E-02 | 8.06 |
|  | GO:0003735~structural constituent of ribosome | 3.23E-02 | 4.03 |
|  | R-MMU-1799339~SRP-dependent co-translational protein targeting to membrane | 6.59E-03 | 5.36 |
|  | R-MMU-975956~Nonsense Mediated Decay (NMD) independent of the Exon Junction Complex (EJC) | 6.59E-03 | 5.25 |
|  | R-MMU-72689~Formation of a pool of free 40S subunits | 9.20E-03 | 4.88 |
|  | R-MMU-156827~L13a-mediated translational silencing of Ceruloplasmin expression | 1.43E-02 | 4.44 |
|  | R-MMU-72706~GTP hydrolysis and joining of the 60S ribosomal subunit | 1.43E-02 | 4.40 |
|  | R-MMU-927802~Nonsense-Mediated Decay (NMD) | 1.43E-02 | 4.36 |
|  | R-MMU-975957~Nonsense Mediated Decay (NMD) enhanced by the Exon Junction Complex (EJC) | 1.43E-02 | 4.36 |
|  | R-MMU-72613~Eukaryotic Translation Initiation | 1.85E-02 | 4.14 |
|  | R-MMU-72737~Cap-dependent Translation Initiation | 1.85E-02 | 4.14 |
|  | KW-0689~Ribosomal protein | 2.12E-02 | 3.80 |
|  | KW-0687~Ribonucleoprotein | 4.78E-02 | 2.81 |
| ***cilium assembly*** |  |  |  |
|  | GO:1905515~non-motile cilium assembly | 5.04E-03 | 9.22 |
|  | GO:0060271~cilium assembly | 1.61E-02 | 4.01 |
|  | GO:0030030~cell projection organization | 4.98E-02 | 4.00 |
|  | KW-0970~Cilium biogenesis/degradation | 9.15E-03 | 3.54 |
| ***Others*** |  |  |  |
|  | GO:0005515~protein binding | 1.09E-04 | 1.58 |
|  | GO:0010273~detoxification of copper ion | 1.31E-02 | 42.8 |
|  | GO:0007623~circadian rhythm | 1.37E-02 | 6.36 |
|  | GO:0001738~morphogenesis of a polarized epithelium | 4.20E-02 | 27.2 |
|  | GO:0007219~Notch signaling pathway | 4.49E-02 | 5.11 |

**Figure 2.**
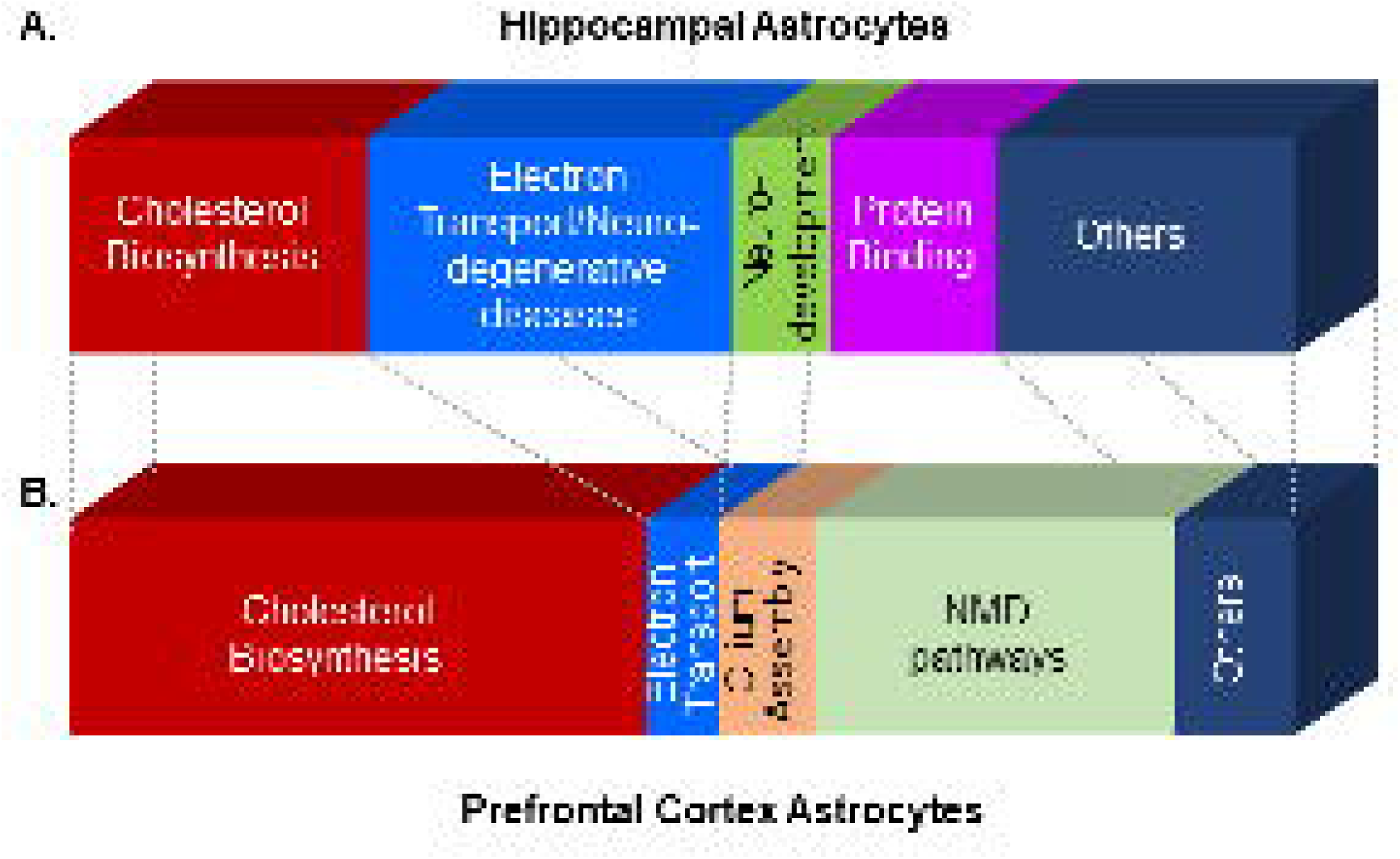
Astrocytes in hippocampus and prefrontal cortex response differently to acute SD. **(A and B)** Biological terms that were enriched in response to acute sleep deprivation were clustered by functional similarity and fractions for the categorized clusters are shown in (**A**) for hippocampal astrocytes and in (**B**) for prefrontal cortical astrocytes. The biological terms that were not clustered are listed as others here and shown in Table 1.

Three common biological networks were identified in astrocytes from both brain regions that responded to acute SD. Among them, clusters of terms related to cholesterol biosynthesis and the mitochondrial electron transport chain were significantly enriched, representing 24.3% and 29.7% of total terms in the HPC, and 47.1% and 5.9% in the PFC, respectively (Figure 2). The circadian rhythm pathway was also enriched in both regions, showing 7.76-fold enrichment in the HPC and 6.36-fold in the PFC (categorized under “others” in Figure 2 and Figure S5A). Beyond these shared pathways, region-specific biological networks were observed: neurodevelopment (Figure 2, Figure S5B) and protein binding (Figure 2 and Figure S5C) pathways were enriched in HPC astrocytes, whereas primary cilia assembly and NMD (nonsense-mediated decay) pathways were unique to PFC astrocytes (Figure 2 and Figure S5D and S5E). Together, these findings suggest that acute SD elicits region-specific molecular responses in astrocytes from the HPC and PFC.

### Cholesterol synthesis pathways were downregulated in both hippocampal (HPC) and prefrontal cortical (PFC) astrocytes following acute sleep deprivation (SD)

As mentioned above, a similarity between astrocytes from the two brain regions was observed in cholesterol metabolism pathways (Figure 2A and 2B). A total of 9 pathway terms and 6 DEGs overlapped between HPC and PFC astrocytes (Figure S6A and S6B). To visualize these results, pathways from the UniProt keyword database were plotted in Sankey diagrams along with enrichment values for HPC (Figure 3A) and PFC (Figure 3B), representing the cholesterol synthesis–associated cluster. The impact of acute SD on cholesterol biosynthesis was more pronounced in the PFC, where a broader range of cholesterol-related pathways were enriched, and a greater number of genes associated with these pathways were differentially expressed.

**Figure 3.**
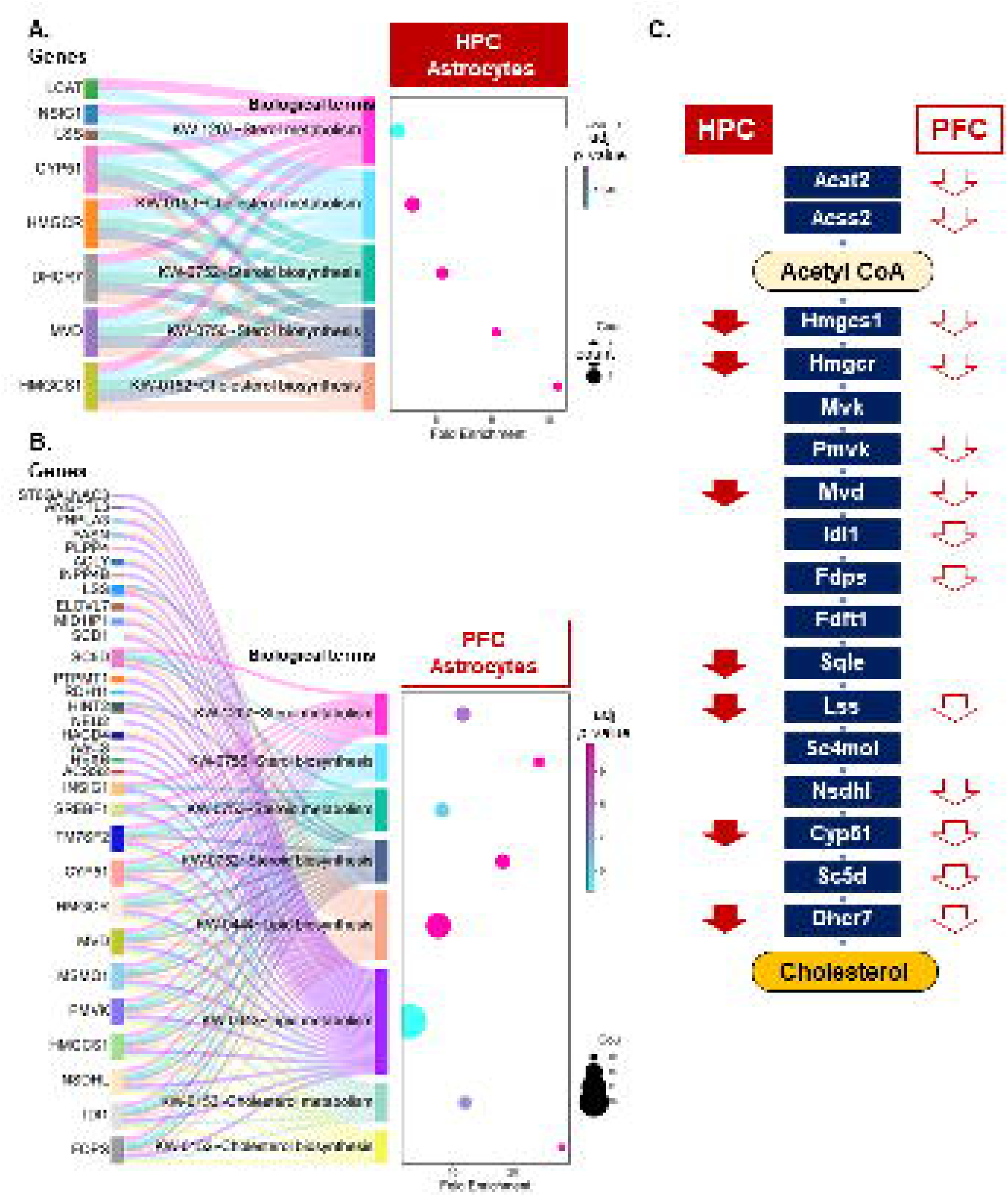
The genes encoding cholesterol synthesis pathways were down regulated in both HPC and PFC astrocytes by acute sleep deprivation. **(A and B)** Sankey diagram (Left) is presented to show associations of the DEGs to the identified terms, along with the enrichment values as a Dot plot (Right). From the total lists of the enriched terms shown in Table 1, the UniProt terms are presented in **(A)** and **(B)** for HPC and PFC, respectively. **(C)** Genes encoding enzymes constituting the de novo cholesterol synthesis pathway were down regulated in astrocytes of HPC or PFC as indicated by down-forwarded arrow. Figure 4. Acute sleep deprivation altered electron transport system in mitochondria in astrocytes of HPC and PFC in opposite directions (A, B and C) and enhanced primary cilia formation in cortical astrocytes (D). Sankey diagram (Left) shows associations of the DEGs to “mitochondrial electron transport/neurodegeneration” related terms, along with the enrichment values shown as a Dot plot (Right) for HPC (A) and PFC (B). (C) The genes contributing enrichment of “Electron transport/Respiratory chain” pathways were altered to opposite direction in HPC (left) and FPC (right) in response to acute sleep deprivation. (D) The intraflagellar transport complex (ITF)-B in astrocytes from PFC was activated following acute SD. The genes listed in the boxes are components of each complex and highlighted in red are up-regulated genes.

Overall, 41 DEGs in cortical astrocytes were assigned to cholesterol biosynthesis–related pathways, accounting for 8.4% of total PFC DEGs—a sharp contrast to the 1.69% of DEGs associated with these pathways in the hippocampus. The total numbers of DEGs were comparable between the two regions (532 in HPC and 486 in PFC; Supplemental Data), suggesting that cholesterol homeostasis in PFC astrocytes may be more sensitive to acute SD than in HPC astrocytes.

In the hippocampus, SD-responsive DEGs included seven genes encoding components of the de novo cholesterol synthesis pathway (Figure 3C), among them *Hmgcr*, which encodes the rate-limiting enzyme 3-hydroxy-3-methylglutaryl-CoA reductase. All seven genes were downregulated by 1.21- to 1.47-fold in SD compared with non-sleep-deprived (NSD) animals (Supplemental Data). Another SD-downregulated gene, *Insig1* (insulin-induced gene 1), regulates cholesterol synthesis in two ways: it inhibits activation of sterol regulatory element-binding proteins (SREBP) - positive regulators of *Hmgcr* transcription - and promotes *Hmgcr* protein degradation by binding to its sterol-sensing domain. In contrast, *Lcat,* which encodes lecithin cholesterol acyltransferase - a plasma enzyme that esterifies cholesterol to facilitate its removal from the cell surface - was the only upregulated gene in the hippocampus following SD.

Similarly, in the PFC, 13 genes involved in de novo cholesterol synthesis were all downregulated by 1.2–1.5-fold (Figure 3C, Supplemental Data). Although most PFC astrocyte DEGs associated with cholesterol synthesis were downregulated (Figure S6B), seven genes were upregulated, including *Slco1b2* (solute carrier organic anion transporter family member 2B1), also known as *Oatp1b2*. In the liver, this transporter facilitates bile acid uptake—a key step in cholesterol excretion—though its role in the brain remains poorly understood. Consistent with these transcriptomic findings, genes encoding two key enzymes in the de novo cholesterol synthesis pathway, *Hmgcr* and *Hmgcs1*, were confirmed to be predominantly expressed in astrocytes (Figure S6C).

Protein interaction analyses using the STRING database further supported these results. The 9 DEGs from HPC astrocytes formed a functionally coherent network related to cholesterol synthesis (Figure S6D), while the 41 DEGs from PFC astrocytes displayed an even denser interaction network. Associations in STRING were based on multiple types of evidence—including text mining, experimental data, database annotations, co-expression, and genomic neighborhood—and are color-coded in Figure S6D.

For further confirmation, additional DAVID analyses were performed on the 74 overlapping DEGs from HPC and PFC astrocytes (Figure S4A) against the UniProt keyword database. A total of 7 cholesterol metabolism related pathways were again returned to be enriched 12.5-fold (Lipid biosynthesis) ∼ 68.3-fold (Cholesterol biosynthesis) as shown in Figure S6E.

### Acute sleep deprivation altered the expression of genes encoding components of the electron transport system in mitochondria in astrocytes of HPC and PFC

The expression of genes involved in the mitochondrial electron transport system was altered in astrocytes from both the HPC and PFC by acute SD (Figure 2A and 2B; Figure 4A and 2B; Figure S7A and S7B). In addition, KEGG pathway analysis revealed that pathways associated with several neurodegenerative diseases were significantly affected in hippocampal astrocytes (Table 1; Figure S7A, B). Neurodegeneration ultimately leads to dysfunction and death of both neuronal and non-neuronal cells through multiple biological processes^33^.

**Figure 4.**
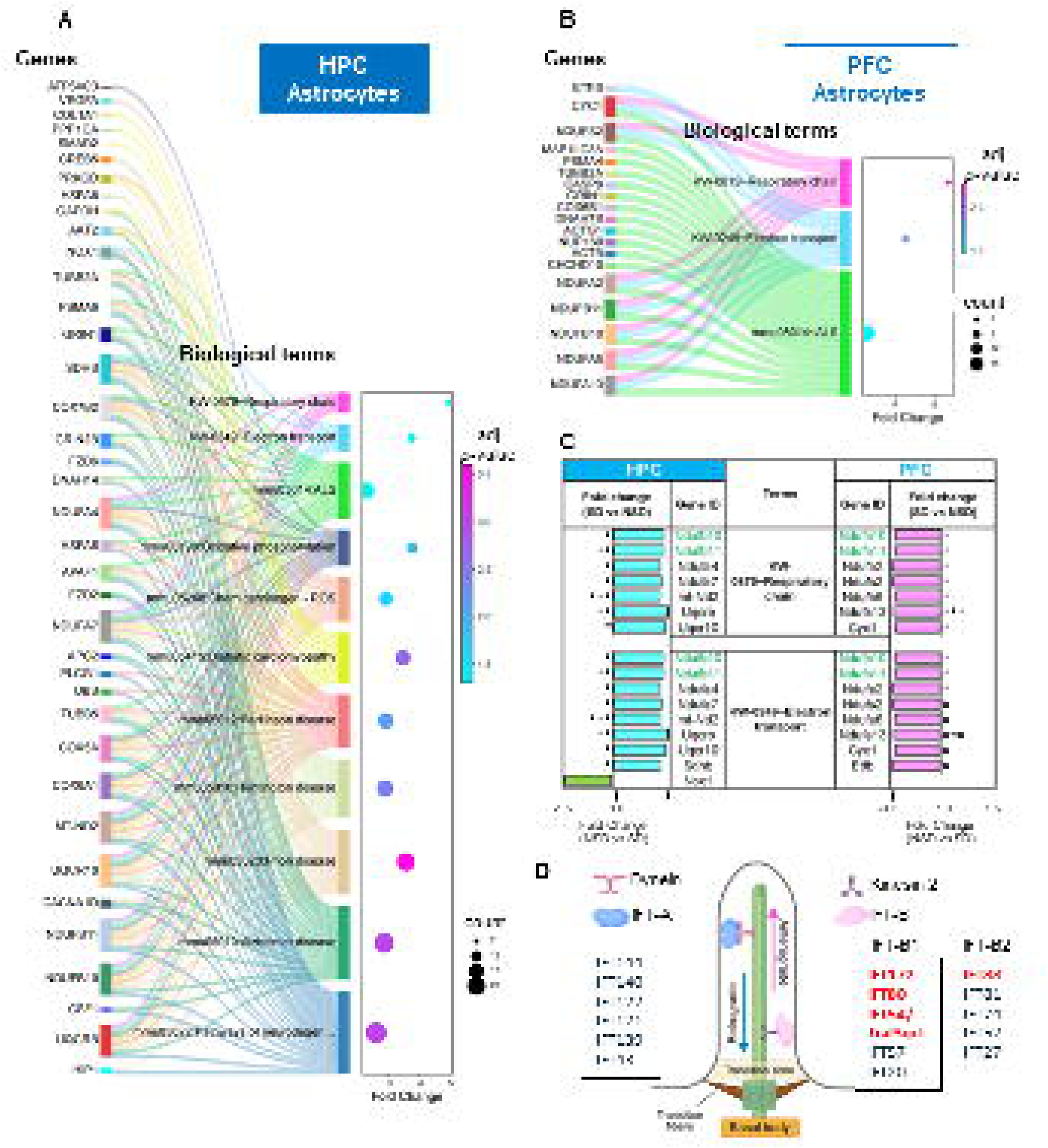
Acute SD altered the genes encoding mitochondria electron transport system, as well as the genes encoding primary cilia assembly in astrocytes. **(A and B)** Sankey diagram (Left) shows associations of the DEGs to “mitochondrial electron transport/neurodegeneration” related terms, along with the enrichment values shown as a Dot plot (Right) for HPC **(A)** and PFC **(B)**. **(C)** The genes contributing enrichment of “Electron transport/Respiratory chain” pathways were altered to opposite direction in HPC (left) and FPC (right) in response to acute sleep deprivation. **(D)** The genes encoding genes involved in primary cilia assembly were modified by acute SD in prefrontal cortical astrocytes, which include those encode the intraflagellar transport complex (ITF)-B. The genes listed in the boxes are known components of each complex and the genes highlighted in red are found to be upregulated in this study. The total DEGs annotated to the primary cilia assembly are listed in Figure S5E.

Functional annotation of the thirty-eight genes associated with these terms indicated that acute SD broadly impacted various neurodegeneration-related processes, with the most pronounced effects on mitochondrial function (Figure S7C, D). This gene cluster included components encoding all classes of the mitochondrial electron transport complexes (I–IV). Although this alternation in the mitochondrial ETC system was observed in the astrocytes from both regions, the direction of these changes was opposite (Figure 4C), indicating that sleep loss causes brain region-specific changes in gene expression. This observation aligns well with the previous studies which exhibited unique transcriptomic profiles in different brain regions following acute SD using the spatial transcriptomic study^34^ and the current study suggest the mitochondria function may be one of those biological pathway that acts differently across the brain region. Among the 74 DEGs shared between HPC and PFC astrocytes, 64 were altered in the same direction, including those involved in cholesterol synthesis (Figure S4A). The rest of 7 genes - including 4 mitochondrial genes - were upregulated in HPC but downregulated in PFC (Figure S4A), whereas 3 genes were vice versa. Interaction analysis using the STRING database confirmed that the thirty-eight DEGs from hippocampal astrocytes formed a functionally connected network, whereas the nineteen DEGs from cortical astrocytes were organized into two smaller, separate networks (Figure S7E).

### Genes involved in primary cilia assembly are upregulated by acute SD

Genes encoding the components of assembly of primary cilia were up regulated in response to acute SD in PFC astrocytes. Non-motile or primary cilia are classified separately from motile cilia which line along epithelial cells and control fluid movement within brain, such as cerebrospinal fluid (CSF). A total of 16 genes were annotated to the 4 terms related to cilium assembly (Table 1 and Figure S5D). Enrichment fold score was highest in the term GO:1905515∼non-motile cilium assembly at 9.2-fold enrichment and the DEGs annotated to the terms were mostly up-regulated (Figure S5D). Primary cilia are built on the basal body, which is originated from a mature mother cell centriole, through multiple steps^35^. It is dynamic organelle that changes its length depending on the environment. Among of the genes aligned to this category, 4 genes belong to the intraflagellar transport (IFT) complex, which moves the essential cilial component back and forth inside cilia along the axoneme. There are two types of IFTs, IFT-A and IFT-B, which act in opposite function, i.e. IFT-A acting as retrograde IFT whereas IFT-B as anterograde IFT, driven by a motor protein, dynein for IFT-A and kinesin 2 for IFT-2 complex. The 4 genes are components of IFT-B and up-regulated, indicating that acute SD promote elongation of primary cilia. Dysfunction of non-motile cilia are known to cause developmental abnormalities in nervous system^36^, more interestingly, its involvement in regulating circadian rhythms and sleep have been reported in multiple studies^37–41^ including its role in maintaining suprachiasmatic nucleus (SCN) network synchrony and circadian rhythms in a Hedgehog signaling manner^37^.

### Down regulation of genes involved in Nonsense Mediated Decay (NMD) in PFC astrocytes by acute SD

There were 15 DEGs that were mapped to 15 biological terms that are associated with Nonsense-mediated Decay (NMD) or translation initiation processes (Figure S5E) in cortical astrocytes. 14 of the DEGs encode ribosomal proteins, 11 of which belong to the 3 NMD related terms from the Reactome (Table 1), suggesting that acute SD modulate the cellular quality control pathway that degrades mRNAs with premature stop-codons to avoid producing truncated proteins^42^, which can potentially interfere with proper protein function in a dominant negative or gain-of-function manner. All DEGs that were assigned to the NMD terms were downregulated and were genes encoding ribosome-binding proteins or small nuclear ribonucleoprotein polypeptide, with an exception of *Ubb*, some of which were found in the 3’-ends of NMD decay intermediates (Figure S5E)^43^.

Our bulk RNAseq analyses on mice hippocampus after acute SD found that *Upf2*, the gene that encodes a scaffold protein to bring the NMD complex together, was up regulated^44^. We did not find the genes encoding Upf2 nor other similar proteins such as Upf1 or Upf3 in the DEG lists from the current study, possibly because MND occurs mainly in cytosol and our analyses was on RNAs originated in nuclear. It requires further investigations to evaluate the possibility that the NMD mechanisms themselves are altered by acute SD.

### Effects of acute sleep deprivation on mature oligodendrocytes (MOLs)

MOLs constituted 32.7% and 17.5% of the total non-neuronal cells in HPC and PFC, respectively, making them one of the largest populations among the cell types identified in our study. We observed significant changes in the transcriptional profiles of MOLs in response to acute SD in both brain regions (Figure S8A). There were 392 and 113 genes that were down- or up-regulated, respectively, in hippocampal MOLs (Figure S8B, Supplemental data). The DAVID analyses using the 505 DEGs from hippocampal MOLs identified 27 biological terms which were further clustered into four categories (Table 2 and Figure S9).

**Table 2.** Enhanced biological pathways in Mature Oligodendrocytes

|  | *Term* | **Benjamini**<br>**adjusted**<br>**p-value** | **Fold**<br>**Enrichment** |
| --- | --- | --- | --- |
| **Hippocampal MOL** |  |  |  |
|  | ***Actin binding*** |  |  |
|  | GO:0003779~actin binding | 7.30E-04 | 3.37 |
|  | GO:0051015~actin filament binding | 4.08E-03 | 3.90 |
|  | KW-0009~Actin-binding | 3.25E-02 | 2.81 |
|  | ***Cell adhesion*** |  |  |
|  | GO:0007155~cell adhesion | 2.73E-04 | 3.13 |
|  | KW-0130~Cell adhesion | 3.07E-02 | 2.28 |
|  | ***Protein folding*** |  |  |
|  | GO:0006457~protein folding | 4.04E-10 | 10.76 |
|  | GO:0140662~ATP-dependent protein folding chaperone | 1.23E-07 | 21.93 |
|  | GO:0051082~unfolded protein binding | 2.39E-06 | 9.19 |
|  | GO:0044183~protein folding chaperone | 1.25E-04 | 11.17 |
|  | GO:0016887~ATP hydrolysis activity | 2.09E-03 | 3.10 |
|  | mmu04141: Protein processing in endoplasmic reticulum | 1.26E-04 | 4.74 |
|  | KW-0143~Chaperone | 3.20E-06 | 4.71 |
|  | KW-0346~Stress response | 5.63E-05 | 7.36 |
|  | ***Protein binding*** |  |  |
|  | GO:0005515~protein binding | 1.23E-07 | 1.65 |
|  | GO:0042802~identical protein binding | 5.72E-03 | 1.73 |
|  | GO:0031625~ubiquitin protein ligase binding | 1.30E-02 | 3.07 |
|  | GO:0031072~heat shock protein binding | 4.04E-02 | 5.98 |
|  | ***Others*** |  |  |
|  | GO:0000166~nucleotide binding | 2.22E-06 | 2.24 |
|  | GO:0004143~ATP-dependent diacylglycerol kinase activity | 2.31E-02 | 23.92 |
|  | GO:0005509~calcium ion binding | 3.47E-02 | 2.16 |
|  | GO:0005524~ATP binding | 1.66E-05 | 2.22 |
|  | GO:0016740~transferase activity | 2.73E-03 | 1.84 |
|  | GO:0016787~hydrolase activity | 1.56E-04 | 2.06 |
|  | GO:0017124~SH3 domain binding | 3.47E-02 | 4.48 |
|  | GO:0031267~small GTPase binding | 3.24E-02 | 3.05 |
|  | GO:0046872~metal ion binding | 4.94E-09 | 1.93 |
|  | mmu05412: Arrhythmogenic right ventricular cardiomyopathy | 4.32E-02 | 5.11 |
| **Prefrontal Cortex MOL** |  |  |  |
|  | ***Protein binding*** |  |  |
|  | GO:0005515~protein binding | 4.22E-03 | 1.65 |
|  | GO:0044877~protein-containing complex binding | 2.43E-02 | 3.17 |

The biggest cluster was a group of pathways related to protein folding that consisted of 43 DEGs (Figure S9A). Those included eight genes encoding heat shock and related proteins, such as *DnaJ heat shock protein family (Hsp40) member A1* (*Dnaja1*) and *A3 (Dnaja3)*, (Figure S9A), all of which were up regulated except *Dnaja3*. Other up-regulated genes include three genes encoding chaperonin subunits, *Cct2*, *Cct3* and *Cct5*, which play a role in autophagic degradation of protein aggregates^45^ (Figure S9A). Three terms related to “actin binding” were also found to be enriched by 2.8 – 3.9-fold in DEGs in hippocampal MOLs (Figure 5A and 5B). The majority of DEGs belong to the terms were up-regulated (Figure S9B).

**Figure 5.**
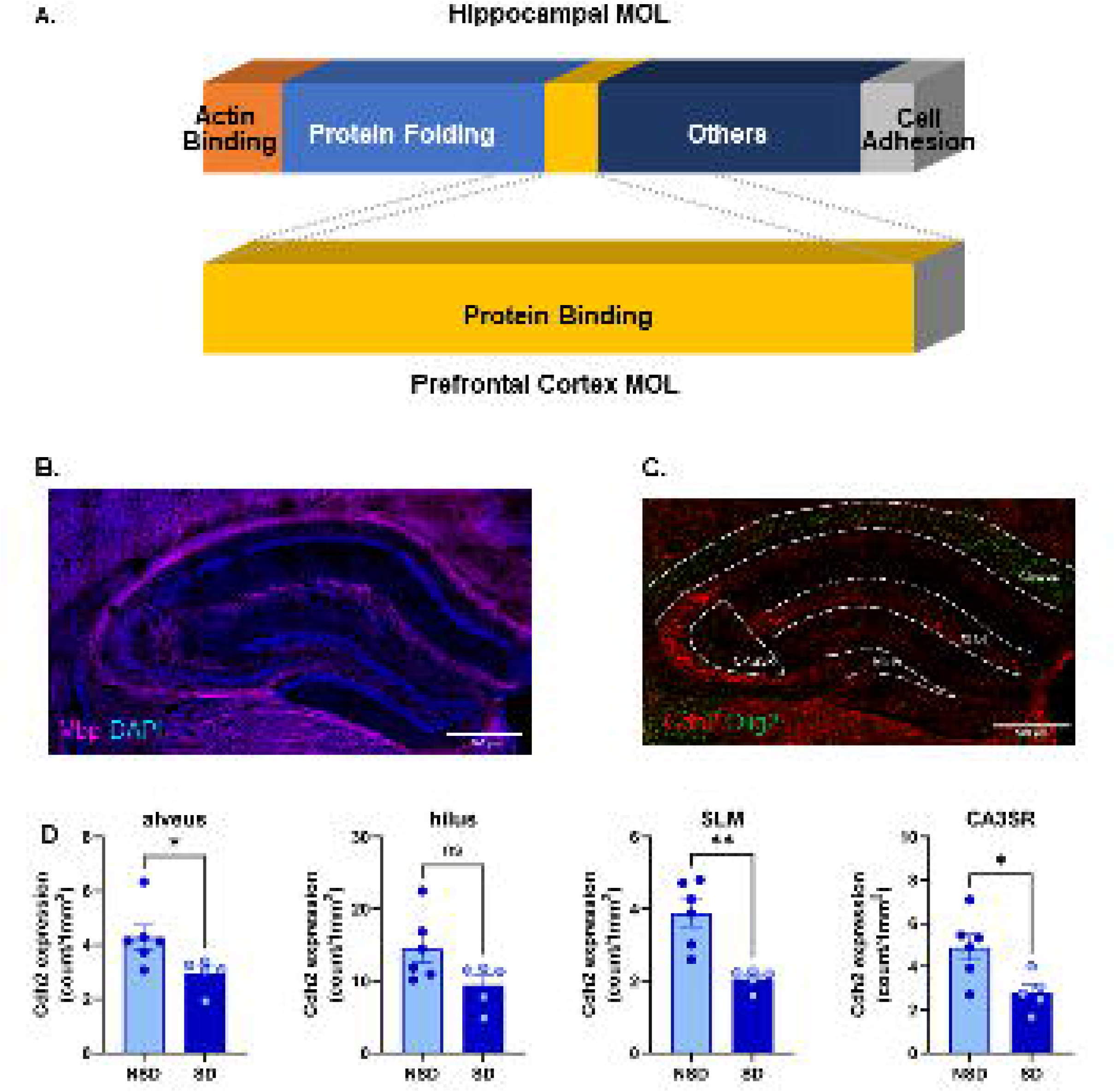
Acute SD impacts distinct biological pathways in mature oligodendrocytes (MOLs). **(A)** Fold enrichment values for the biological terms are shown as bar graph and clustered based on functional similarity. The color of the bars indicates Benjamini’s adjusted p-values. **(B)** Fractions of the clusters of pathways that were enriched by scute sleep deprivation identified in HPC (top panel) and PFC (bottom panel). “Protein folding” was the biological term that was the most significantly enriched in hippocampal MOL. **(C)** Antibody for Myelin basic protein (MBP) was used to localize matured oligodendrocytes (MOL). **(D)** Sections containing hippocampus were analyzed for expression level of *Cadherin2* in oligodendrocytes regions. An Oligodendrocyte marker, Olig2 was used to localize oligodendrocytes. **(E)** RNAscope analyses confirmed the downregulation of Cdh2 in alveus, SLM and CA3SR. Unpaired two-tailed t-tests were used to assess the significance of Cdh2 expression level changes between NSD (N=6) and SD (N=5). *: p<0.05, **: p<0.01, ns: not significant. Representative RNAscope images that were used for the analyses are shown in Figure S7B.

Another interesting biological process that was affected by acute SD was functions that are affected by cell adhesion molecules. Expression of cell adhesion proteins encoded by the genes listed in Figure S9C were altered in both directions in HPC MOLs. We observed 29 genes encoding adhesion proteins which span multiple adhesion subfamilies such as Ca^2+^-independent, or -dependent, Immunoglobulin superfamily CAMs (IgSF CAMs), and integrins. 6 of them were up regulated and 23 down regulated, with no apparent association to a certain subfamily (Figure S9C).

In cortical MOLs, 119 up-regulated and 150 down-regulated DEGs were identified (Figure S8B). However, only two terms that were related to “protein binding” passed Benjamini’s adjusted p-value of 0.05 on DAVID analyses using the total 269 genes (Table 2). With the use of the currently available databases, it was difficult to identify significant functions that were affected by acute sleep deprivation in cortical MOLs.

### Sleep deprivation down-regulates Cdh2 expression in MOLs

Adhesion molecules are vital for preserving the structural integrity of the myelinated unit, which is required for rapid saltatory conduction. They facilitate the proper wrapping of myelin sheath around the axon by oligodendrocytes in the central nervous system^46^. As mentioned above, we saw a number of adhesion molecules responded to acute SD in MOL of HPC. Among these genes, the neural-cadherin (*Ncad*) gene *Cdh2* was the most down regulated in MOL originated from HPC (Figure S9A). *Ncad* is a classical cadherin protein that mediates cell-cell interaction in a Ca2+-dependent manner in the central nervous system^47^, and is involved in axon-oligodendrocyte contact and myelination^48^.

To confirm the SD-induced reduction of Cdh2 and identify the most affected hippocampal sub-regions, we performed RNAscope experiments targeting *Cdh2* and *Olig2* (a reference marker for oligodendrocytes) (Figure 5B and 5C). Based on the *Olig2* and *Mbp* (a myelination marker protein) expression patterns, we selected the alveus, stratum lacunosum-moleculare (SLM), hilus within the dentate gyrus, and stratum radiatum close to CA3 (CA3SR) as target regions (Figure 5C). We compared the *Cdh2* mRNA expression levels in each region using multiple approaches including fluorescence intensity and cell count measurements The results indicated a significant 1.5 to 2-fold reduction of *Cdh2* mRNA in alveus, SLM and CA3SR after SD. Although not significant, *Cdh2* mRNA also showed a trend for reduction within the hilus (Figure 5D).

The SLM is a critical interface where CA1 pyramidal neurons receive synaptic inputs from the entorhinal cortex (EC) via the temporoammonic (TA) pathway^49^. The SLM also includes inhibitory interneurons that coordinate EC and CA3 dual inputs to CA1, thereby switching the hippocampus between encoding and retrieval modes^50^. Given the role of *Cdh2* in neuron-oligo contact and myelination^48,51^, the reduction in the Cdh2 mRNA level, especially in the SLM, might indicate deficits in hippocampal neuronal axon myelination, altered synaptic transmission and plasticity. Such changes could impair information processing both within the hippocampus and between the hippocampus and other brain regions.

## DISCUSSION

Sleep has traditionally regarded as a neural process, yet increasing evidence indicates that astrocytes and other non-neuronal cells play essential role in maintaining brain homeostasis during the sleep–wake cycle^17,52,53^. Despite this recognition, the molecular responses of these cells to sleep loss remains poorly understood. In this study, we used single-nucleus transcriptomics for enriched non-neuronal cells—focusing on astrocytes and MOLs—to examine the effects of acute sleep deprivation (SD) in two brain regions central to cognition: the HPC and PFC. We demonstrate that acute SD elicits robust, cell type- and region-specific transcriptional reprogramming. The most prominent effects observed are the widespread suppression of cholesterol biosynthesis genes in astrocytes across both regions studied. There is a notable, regionally opposing regulation of mitochondrial and neurodegeneration-related pathways. Additionally, genes associated with primary cilia are induced in cortical astrocytes, while genes encoding cell adhesion molecules, including Cdh2 (N-cadherin), are down-regulated in oligodendrocytes. These findings provide molecular evidence for previously suggested phenotypes and identify the specific cell types responsible for them^54^ and complemented the findings came from the similar studies on neuronal cells^44,55–59^. Together, these results highlight the role of non-neuronal cells in the brain’s response to sleep loss, expanding the traditional neuron-centric view of sleep regulation.

### Reduced cholesterol and cell adhesion molecules as possible contributors to response acute SD

Astrocytes play pivotal roles in maintaining the brain’s metabolic balance by regulating ion homeostasis, supplying energy substrates, and synthesizing lipids essential for synaptic remodeling^60,61^. Our data show that acute SD reduces the expression of genes in the cholesterol biosynthetic pathway, including *Hmgcr*, *Hmgcs1*, *Mvd*, and *Insig1*, which together coordinate SREBP-dependent cholesterol synthesis. Astrocyte-derived cholesterol is required for neuronal membrane renewal, dendritic spine formation, neurotransmitter receptor localization and myelin formation^62,63^.

Myelin, the insulating sheath surrounding axons made in oligodendrocytes, enhances the speed and efficiency of nerve impulse conduction^64^ and cholesterol is one of its major structural component. During developmental stages, oligodendrocytes can synthesize majority of cholesterol supply on its own, however, astrocytes become a more significant source of cholesterol in adults^65^. The astrocytes produce cholesterol and package it into lipoproteins, primarily those containing ApoE to deliver to other cells. Therefore, reduced cholesterol synthesis in astrocytes during SD could negatively affect myelin formation. Myelin plasticity has recently been recognized as an important modulator of neuronal network activity, including circuits involved in sleep homeostasis^66^.

Beyond myelination, cholesterol is also a key component of cellular membranes, influencing membrane fluidity and the conformation of membrane proteins such as ion channels and receptors^67,68^. Thus, altered cholesterol levels may contribute to the reduction in hippocampal synaptic plasticity observed after acute SD^69,70^. Non-neuronal cells are increasingly recognized for their roles in regulating synaptic transmission and plasticity through neuron-glia communication^71–73^. Multiple studies have shown that astrocyte-derived cholesterol promotes synaptogenesis^74–76^ and may enhances synaptic stability^77,78^. Future studies will be needed to explore these ideas.

Additionally, acute SD significantly downregulated a cluster of genes encoding cell adhesion molecules in hippocampal oligodendrocytes (Figure S9C), potentially further destabilizing synaptic plasticity. Among these, *Cdh2* was markedly reduced in a tissue-specific manner. RNAscope validation confirmed decreased *Cdh2* mRNA in hippocampal regions rich in myelinated axons and entorhinal inputs, including the alveus and stratum lacunosum-moleculare (SLM). Because N-cadherin-mediated adhesion is essential for activity-dependent myelination^79^, reduced *Cdh2* expression likely reflects weakened axon–glia coupling and disrupted conduction efficiency.

### Cholesterol dysregulation and neurodegenerative pathologies

The bidirectional relationship between sleep dysregulation and neurodegenerative pathology is well documented in both clinical and experimental studies^80–86^. Although the correlation between sleep loss and progression of neurodegenerative disease is strong, the underlying mechanisms remain incompletely understood. Our finding that acute SD suppresses cholesterol biosynthesis suggests a potential pathway linking sleep loss and neurodegeneration.

In Alzheimer’s disease (AD), astrocyte-derived cholesterol plays a pivotal role in β-amyloid (Aβ) generation^87–89^. Cholesterol synthesized in astrocytes is transported to neuronal membranes, where it supports the formation of lipid rafts—specialized microdomains that promote Aβ cleavage from the amyloid precursor protein (APP)^87^. Consistent with this, metabolomic and transcriptomic studies have revealed reduced cholesterol levels in brain tissues from AD brain^90^. Moreover, a recent meta-analyses of single-nucleus RNA sequencing (snRNA-seq) datasets show down-regulation of lipid metabolic processes in astrocytes across AD stages^91^.

Similar lipid-related abnormalities are implicated in Huntington’s disease (HD) and Parkinson’s disease (PD)^92–101^. In HD, mutant huntingtin (mHTT) disrupts the SREBP2/importin-β complex—a key regulator of lipid metabolism—thereby disrupting cholesterol biosynthesis^92^. In PD, α-synuclein (α-syn) aggregates within Lewy bodies and interacts with membrane lipids, impairing ER-to-Golgi trafficking^93,100,101^. Interestingly, inhibition of HMG-CoA reductase, the rate-limiting enzyme in cholesterol biosynthesis, with lovastatin reduces neurite degeneration in both animal and cellular models of PD^97^. These parallels reinforce the notion that lipid dysregulation and glial metabolic dysfunction are central to both sleep loss and neurodegenerative vulnerability.

### Regional heterogeneous responses to acute sleep deprivation: cortical vulnerability and mitochondrial adaptation

Although cholesterol biosynthetic genes were downregulated in astrocytes of both the HPC and the PFC, the suppression was stronger and more extensive in the PFC. Despite similar total numbers of differentially expressed genes, cholesterol-related pathways were more prominently enriched in the PFC. This asymmetry suggests that cortical astrocytes are particularly sensitive to metabolic stress induced by wakefulness. This finding aligns with evidence that the PFC is among the first regions to exhibit metabolic and functional decline during sleep deprivation^102,103^, likely due to its high energetic demands associated with executive function.

In addition to lipid metabolism, astrocytes displayed marked reorganization of mitochondrial gene networks, with opposing effects between the two regions. In hippocampal astrocytes, genes encoding components of the electron transport chain (ETC) - including genes encoding the enzymes that belong to Complex I such as *Ndufa, Ndufb family*, Complex IV such as *Cox* family such as - were up-regulated, indicating enhanced oxidative phosphorylation. In contrast, a similar set of genes was downregulated in PFC astrocytes, suggesting mitochondrial suppression or stress. This dichotomy mirrors previous observations that hippocampal neurons transiently boost oxidative metabolism during wakefulness, whereas cortical neurons accumulate oxidative damage under prolonged activity^104^. Such regional heterogeneity was evident in our spatial transcriptomic analysis using the 10x Genomics Visium platform^34^. and also in the report of non-uniform, sleep-dependent neural reactivation during memory consolidation^105^. The enrichment of neurodegeneration-related pathways in hippocampal astrocytes further implies that transient hypermetabolism may precede oxidative stress. Together, these opposite responses may represent region-specific metabolic strategies—adaptive activation in the hippocampus versus an energy-conserving or protective state in the cortex. Beyond these changes in mitochondral gene networks, cholesterol serves as a structural component of mitochondrial membranes and modulates the efficacy of oxidative phosphorylation^106–108^. Thus, the downregulation of cholesterol biosynthetic genes in astrocytes may indirectly compromise mitochondrial performance, exacerbating energy imbalance during sleep loss. Considering recent findings that mitochondrial-derived electrons may act as sleep-promoting signals^109^, delineating region-specific mitochondrial responses to acute SD will be an important direction for future research.

### Up-regulation of astrocytic primary cilia - potential sensors of sleep pressure

An additional, unexpected finding was the up-regulation of genes associated with primary (non-motile) cilia in PFC astrocytes following SD. Primary cilia act as sensory hubs that integrate extracellular cues—such as metabolic, hormonal, and circadian signals—into transcriptional responses, often in a cell specific manner^110–112^. Primary cilia are well recognized for their roles in developmental, sensory, and homeostatic regulation^36,113^, and increasing evidence links ciliary dysfunction to neurological, psychiatric, and metabolic disorders^114–118^. In the suprachiasmatic nucleus (SCN) of the hypothalamus in the mouse brain, primary cilia undergo oscillations in both density and length on a 24-hour cycle, being most condensed and elongated at ZT0, when sleep needs are elevated^37^. Our study found a force driving primary cilia elongation at ZT5, after 5 hours of SD, which may reflect adaptive mechanisms aimed at maintaining sleep homeostasis. However, since the oscillation of astrocytic primary cilia remains unconfirmed, this hypothesis may be premature. Alternatively, the activation of primary cilia assembly following acute SD might indicate the initiation of neurotoxic effects. In astrocytes, Muhamad et al. demonstrated that primary cilia regulate Complement C3 production in reactive astrocytes^119^. They observed cilia elongation in C3-positive, neurotoxic astrocytes, while conditional ablation of cilia reduced astrocyte activation and C3 expression^119^. Since C3 activation can trigger cell death, enhanced ciliogenesis may represent an additional mechanism through which acute SD promotes astrocytic or neuronal apoptosis, complementing mitochondrial-mediated pathways^120^.

Adding more complexity to understand the roles of astrocytic primary cilia in sleep regulation, the concurrent down-regulation of cholesterol biosynthetic genes and induction of ciliary genes has to be addressed given that membrane cholesterol influences ciliary receptor localization and membrane fluidity^121^, reduced cholesterol synthesis may compromise ciliary structure or signaling capacity. Ciliary remodeling may thus serve as a critical link between glial metabolism and the circadian and behavioral regulation of sleep.

### Integrated model of glial adaptation to sleep loss

Overall, our findings support a model in which acute SD induces coordinated yet regionally divergent transcriptional reprogramming across glial populations. In astrocytes, suppression of cholesterol synthesis and mitochondrial remodeling constrain energy-intensive processes, while up-regulation of ciliary genes may recalibrate responsiveness to extracellular cues. In oligodendrocytes, decreased cell adhesion activity and altered cytoskeletal and chaperone gene expression indicate weakened axon–glia coupling and increased cellular stress. Together, these adaptations may transiently reduce glial support for neurons, contributing to the cognitive and emotional deficits observed after even brief periods of sleep loss.

Importantly, the astrocytic and oligodendrocytic responses appear interconnected. Reduced cholesterol biosynthesis may impair both neuronal membrane function and myelin maintenance, as cholesterol is a major lipid component of myelin. Similarly, mitochondrial dysfunction in astrocytes could limit the energy substrates available to oligodendrocytes, further compromising myelin stability. These cross-cellular effects emphasize the importance of glial coordination in maintaining neural circuit integrity during sleep–wake transitions. By identifying the glial pathways most sensitive to sleep loss, our findings underscore the importance of non-neuronal cells in sustaining brain metabolic homeostasis and neural connectivity, offering new avenues for understanding and mitigating the cognitive consequences of sleep deprivation.

## LIMITATIONS AND FUTURE DIRECTIONS

While this study provides a comprehensive transcriptomic profile of non-neuronal responses to acute sleep loss, several limitations should be acknowledged. First, all analyses were conducted at a single post-deprivation time point, which precludes assessment of molecular and cellular recovery during rebound sleep. Longitudinal studies integrating time-resolved transcriptomics, lipidomic, and metabolomic analyses will be necessary to determine whether these molecular changes are transient adaptations or predictors of chronic dysfunction. Second, although strong correlations were observed between gene expression changes and metabolic pathways, functional validation at the protein and physiological levels remains essential. Direct measurement of cholesterol and ATP content, mitochondrial activity and states, and myelin structure, will help clarify the biological consequences of these transcriptional alterations. Finally, integrating neuronal data with glial transcriptomics and spatial mapping could provide deeper insight into how non-neuronal reprogramming influences neural circuit dynamics during sleep deprivation.

## RESOURCE AVAILABILITY

### Lead contact

Further information and requests for resources and reagents should be directed to and will be fulfilled by the lead contact, Ted Abel.

### Materials availability

This study did not generate new unique reagents.

### Data and code availability

- Single-nuclei RNA-seq data have been deposited at GEO: accession number is GSE313497 and is publicly available as of the date of publication.
- Any additional information required to reanalyze the data reported in this paper is available from the lead contact upon request.

## ACKNOWLEDGMENTS

We thank Deaven Denis, Jacob Belardo, Taylor Voyna and Tasha Gilkison (University of Iowa) for their technical assistance with genotyping. JK appreciates Dr. Toshi Kitamoto (University of Iowa) for the critical reading of the manuscript and the constructive advice.

The data presented herein were obtained at the Flow Cytometry Facility, which is a Carver College of Medicine / Holden Comprehensive Cancer Center core research facility at the University of Iowa. The facility is funded through user fees and the generous financial support of the Carver College of Medicine, Holden Comprehensive Cancer Center, and Iowa City Veteran’s Administration Medical Center.

The data presented herein were also obtained at the Genomics Division of the Iowa Institute of Human Genetics which is supported, in part, by the University of Iowa Carver College of Medicine and the Holden Comprehensive Cancer Center (National Cancer Institute of the National Institutes of Health under Award Number P30CA086862).

## AUTHOR CONTRIBUTIONS

Conceptualization, J.K., M.F., and T.A.; Methodology, J.K. Y.V. Y.W. and T.A.; Investigation, J.K., M.F., and T.A.; Writing - original draft, J.K.; Writing - review & editing, J.K., Y.W., M.F., and T.A.; Funding acquisition, M.F., and T.A.; Resources, M.F., and T.A; Supervision, M.F., and T.A.

## DECLARATION OF GENERATIVE AI AND AI-ASSISTED TECHNOLOGIES

Not Applicable.

## SUPPLEMENTAL INFORMATION

Supplemental Figures: S1–S10.

Supplemental Figures Legends: Word file containing Supplemental Figure Legends.

Supplemental Data: Excel file containing 1) Parameter used in snRNAseq analyses 2) List of cell markers, 3) Summary of the numbers of differentially expressed genes (DEGs) for astrocytes and MOLs in HPC and PFC, 4) lists of differentially expressed genes (DEGs) for astrocytes, 5) and for MOLs and 6) transcriptional profiles of sub types of astrocytes and MOLs from HPC and PFC.

## STAR⍰METHODS

### EXPERIMENTAL MODEL AND STUDY PARTICIPANT DETAILS

N/A

### METHOD DETAILS

#### Mice stocks, maintenance and breeding

B6;129-*Gt (ROSA)26Sor^tm^*^5^(CAG–Sun1/sfGFP)*^Nat^*/J (JAX:021039) males and B6; FVB-Tg (Aldh1l1-cre)JD1884Htz/J (JAX:023748) females were obtained from The Jackson Laboratory (Bar Harbor, ME, USA). Hemizygous *Aldh1l1-Cre +/−* female mice were bred with homozygous *CAG-Sun1/sfGFP* +/+ males. Pups that were hemiozygous with the *Aldh1l1-cre* and the *Sun1/sfGFP* transgenes were used for experiments. Mice were housed in standard cages at an ambient temperature of 24 ± 0.5°C on a 12/12 h light/dark cycle with food and water ad libitum. All experimental procedures were approved by the Institutional Animal Care and Use Committee of the University of Iowa and conducted in accordance with National Research Council guidelines and regulations for experiments in live animals.

#### Sleep Deprivation

A pair of 11-12-week-old littermate males with one copy of the *Aldh1l1*-cre and the *Sun1*/sfGFP transgenes were single-housed for five days prior to the experiment day. Each cage had corncob bedding (Envigo, Teklad ¼” corncob, #7907) and a small amount of soft bedding for nest building. These cages were equipped with water bottles and wire hoppers to hold food. Mice had ad libitum access to food and water. While being single housed, the mice were taken to the procedure room once a day and received brief gentle tapping and shaking to acclimate to the sleep deprivation procedure. This allowed the mice to habituate to transportation to the room and to the tapping stimulation.

A mouse of the pair was subjected to an acute sleep deprivation (SD) beginning at ZT0 and continued for 5 hours, while the other mouse was continued to be housed in the home cage. The SD was performed using the gentle handling method, in which the cage was gently tapped as needed to keep the mice awake^122^. When tapping was no longer sufficient, the mice received a “fresh air” by briefly removing the lid to refresh the air inside the cage or a gentle shaking. At ZT5, a cage was moved to the dissection room where the hippocampus or PFC was dissected out, briefly rinsed in PBS, and then flash-frozen in liquid nitrogen. Two or three pairs of mice were used for snRNAseq analyses of hippocampus or prefrontal cortex, respectively. The frozen tissues were kept in a -80°C freezer until use.

#### Immunochemistry

Mice used for immunohistochemical staining (IHC) were perfused with 1× PBS, followed by cold 4% paraformaldehyde (PFA). Brains were dissected into 4% PFA solution and fixed overnight at 4°C before being transferred to 30% sucrose solution and kept for at least 48 hours. Equilibrated brains were sliced into 30 μm-thick sections on a cryostat (Leica 3050S) at -20 °C and kept in cryoprotectant solution (30% Sucrose, 30% Ethylene Glycol, and 0.01% Sodium Azide diluted in 1X PBS). Slices of dorsal hippocampus were washed in PBS and incubated for 2 hours in blocking solution (PBS containing 0.2% Triton X-100 and 3% goat serum). [After rinsing with PBS-T (containing 0.2 % Triton X-100), the slices were incubated with primary antibody (mouse anti-NeuN antibody, Abcam #ab104224) at 1:500 dilution, mouse anti-GFAP (Cell Signaling Technology #3670S) at 1:1000 dilution, rabbit anti-SOX9 antibody (Abcam #ab185966) at 1:500 dilution, and rabbit Recombinant Monoclonal Myelin Basic Protein EPR21188 (Abcam #ab218011) at 1: 5000 in blocking buffer for overnight in a cold room. After another round of washing with PBS-T, the sections were incubated with secondary antibody (Alexa Fluor™ 594 conjugated anti-mouse, or anti-rabbit (Invitrogen #A-11005 or Invitrogen #A11012, respectively) at 1:500 in blocking buffer for 2 hours and followed by a final wash with PBS before mounting on glass slides.

Following IHC, images of dorsal hippocampus were acquired using a Leica SPE Confocal microscope equipped with lasers at 405nm, 488nm, 561nm, and 635nm. All images (8-bit) were obtained with identical settings for laser power, detector gain and pinhole diameter (1.0AU) using 20X or 40X oil immersion objectives at 1024 x 1024-pixel resolution and 1.5X optical zoom.

#### RNAscope

RNAscope in situ hybridization was performed to assess N-Cadherin (Cdh2) expression in oligodendrocytes using the RNAscopeTM Multiplex Fluorescent Reagent Kit v2 (Advanced Cell Diagnostics, Newark, CA, US). Brains were dissected from SD and NSD mice at ZT5 and post-fixed in 4% paraformaldehyde in PBS overnight at 4°C. Brains were transferred stepwise to 10%, 20% and 30% sucrose in PBS and kept at 4°C. In brief, fixed brains were sectioned on a cryostat (16 um) and mounted on Superfrost^TM^ Plus microscope slides (Fisher Scientific, Cat. #12-550-15). Slides were then underwent serial dehydration steps in 50%, 70%, and 100% ethanol, followed by Hydrogen Peroxide (Advanced Cell Diagnostics, #322330) treatment at room temperature. Slides were then pretreated with Target Retrieval and Protease III according to the manufacturer’s instructions. Hybridization of probes against Cdh2 (Cat.489571) was done at 40°C for 2h in an HybEZ oven (Advanced Cell Diagnostics, Newark, CA, US). After probe hybridization, signals were amplified with AMP1, AMP2, and AMP3, and counter stained with OPAL dyes @ 520, 570, and 690 (Akoya Biosciences). Slides were mounted with prolong diamond antifade mountant with DAPI (Thermofisher, Cat. #P369632) to visualize nuclei and stored at 4°C. Images were acquired using a Leica SPE Confocal Microscope with 10X and 63X objectives and processed using ImageJ software.

#### Isolation of nuclei from hippocampus and prefrontal cortex

The nuclei were isolated from frozen hippocampus or PFC using the 10x Genomics Chromium Nuclei Isolation with RNase Inhibitor Kit (cat# 1000494, 10x Genomics, Pleasanton, CA) according to the provided protocol. All procedures were carried out on ice or at 4°C. Briefly, the tissues were first lysed and homogenized in 0.5 mL of lysis buffer in a 1.5 mL microtube using a plastic pestle and incubated on ice for 10 minutes to release the nuclei. The homogenized tissue was then filtered, followed by centrifugation at 500 rcf for 3 minutes. The precipitates were resuspended in 0.5 mL of Debris Removal buffer and spun at 700 rcf for 10 minutes. The precipitates were washed with 1 mL of wash buffer once and resuspended in 1 mL of resuspension buffer. Anti-NeuN antibody (Recombinant Alexa Fluor® 647 Anti-NeuN antibody, cat #190565, Abcam, Waltham, MA) was added to the nuclei suspension at a 1:500 dilution and incubated for 30 minutes in a cold room by gentle agitation. After another wash with 1 mL of wash buffer, the nuclei were resuspended in 300 µL of resuspension buffer containing Hoechst dye at 4 ng/mL. The extracted nuclei were sorted on a BD12 FACSAria Fusion at the Flow Cytometry Facility of the University of Iowa. The optimal gate settings were determined using control nuclei samples with GFP or Alexa Fluor® 647 fluorescent signal alone. Sorting was done on individual nuclei with a 100 um nozzle based on the presence or absence of either GFP or Alexa Fluor® 647 and all four sub-fractions were collected into separate microtubes. The sub-fraction with only GFP signals was stained with AOPI dye (Logos Biosystems, Gyeonggi-do, South Korea, cat # F23001) and the concentration of nuclei was determined using a LUNA-FL™ Dual Fluorescence Cell Counter (Logos Biosystems, Gyeonggi-do, South Korea) to calculate a target number prior to capture. All procedures were done at 4C or on ice.

#### Single nuclei RNAseq

The fraction with GFP-positive nuclei was partitioned into single droplets using the Chromium X (10x Genomics, Pleasanton, CA). cDNA synthesis and single-nuclei RNAseq library construction was carried out using the Chromium GEM-X Single Cell 3’ Library Preparation Kit V3.1 (10x Genomics, Pleasanton, CA) according to the manufacturer’s instructions at the Iowa Institute for Human Genetics at University of Iowa. The amplified cDNA libraries were sequenced on the Illumina NovaSeq6000 Sequencing System with a paired 50 bp run format.

#### Data processing and analyses

Quality assessment and controls (QA/QC), alignment to the genome database, and quantification analyses were performed using the 10x Genomics Cell Ranger for Gene Expression tool. Downstream analyses were conducted using the Partek^TM^ Flow^TM^ software (Build version 11.0.24.0226, Partek/Illumina, San Diego, CA). Briefly, raw FASTQ files were processed through 10x Genomics Cell Ranger for Gene Expression. The returned reads were evaluated for total read counts, the numbers of detected features, mitochondria gene counts and ribosomal RNA counts. After filtering the total reads using the cut off values listed in the supplemental data, the filtered reads were aligned against the Mus musculus mm39 database with the Ensembl Transcript release 105 as the index, and normalized for use in SCT, PCA, UMAP, clustering, and differentially expressed gene (DEG) analyses. Detailed parameters for each task, as well as cell markers that were used to classify cell types are recorded in the supplemental data. Differentially expressed genes in response to an acute sleep deprivation were identified based on normalized values by ANOVA (Analyses of Variance) method for each cell types using a cut off value of adjusted p of 0.05 or less and fold change of more or less than I1.2I.

Biological significance of the DEGs were assessed using The Database for Annotation, Visualization, and Integrated Discovery (DAVID) v2023q4 (<u>DAVID Functional Annotation</u> <u>Bioinformatics Microarray Analysis</u>) against the Gene Ontology, Uniprot Keyword, KEGG Pathway, and REACTOME Pathway databases. The terms with less than 5% of Benjamini-adjusted p-values were considered significant.

### QUANTIFICATION AND STATISTICAL ANALYSIS

All data presented in this study were organized and analyzed on Graph Pad Prizm 10 by the statistical methods indicated in figure legends.

### ADDITIONAL RESOURCES

Please provide links to websites that provide further information relevant to the study (e.g., protocol download, troubleshooting forum, etc.). Clinical trial registry numbers and links should also be placed here. Please briefly describe the resource and its relevance for the paper. Please report this information as: “Description: URL.”

## REFERENCES

1. Colten HR, A.B. (2006). Sleep Disorders and Sleep Deprivation: An Unmet Public Health Problem. In Extent and Health Consequences of Chronic Sleep Loss and Sleep Disorders, (National Academies Press (US)).

2. Reichert, C.F., Deboer, T., and Landolt, H.P. (2022). Adenosine, caffeine, and sleep-wake regulation: state of the science and perspectives. J Sleep Res 31, e13597. 10.1111/jsr.13597.

3. Chua, E.C.-P., Fang, E., and Gooley, J.J. (2017). Effects of total sleep deprivation on divided attention performance. PLOS ONE 12, e0187098. 10.1371/journal.pone.0187098.

4. Dopp, J., Ortega, A., Davie, K., Poovathingal, S., Baz, E.-S., and Liu, S. (2024). Single-cell transcriptomics reveals that glial cells integrate homeostatic and circadian processes to drive sleep–wake cycles. Nature Neuroscience 27, 359–372. 10.1038/s41593-023-01549-4.

5. Knutson, K.L., and Van Cauter, E. (2008). Associations between Sleep Loss and Increased Risk of Obesity and Diabetes. Annals of the New York Academy of Sciences 1129, 287–304. 10.1196/annals.1417.033.

6. Van Cauter, E., Holmback, U., Knutson, K., Leproult, R., Miller, A., Nedeltcheva, A., Pannain, S., Penev, P., Tasali, E., and Spiegel, K. (2007). Impact of sleep and sleep loss on neuroendocrine and metabolic function. Horm Res 67 Suppl 1, 2–9. 10.1159/000097543.

7. García-Marín, V., García-López, P., and Freire, M. (2007). Cajal’s contributions to glia research. Trends in Neurosciences 30, 479–487. 10.1016/j.tins.2007.06.008.

8. Frank, M.G. (2013). Astroglial regulation of sleep homeostasis. Curr Opin Neurobiol 23, 812–818. 10.1016/j.conb.2013.02.009.

9. Frank, M.G. (2016). Shining a light on astrocytes and sleep (Commentary on Pelluru et al.). Eur J Neurosci 43, 1297. 10.1111/ejn.13148.

10. Frank, M.G. (2019). The Role of Glia in Sleep Regulation and Function. Handb Exp Pharmacol 253, 83–96. 10.1007/164_2017_87.

11. Halassa, M.M., Florian, C., Fellin, T., Munoz, J.R., Lee, S.-Y., Abel, T., Haydon, P.G., and Frank, M.G. (2009). Astrocytic Modulation of Sleep Homeostasis and Cognitive Consequences of Sleep Loss. Neuron 61, 213–219. 10.1016/j.neuron.2008.11.024.

12. Andersen, J.V., and Schousboe, A. (2023). Milestone Review: Metabolic dynamics of glutamate and GABA mediated neurotransmission — The essential roles of astrocytes. Journal of Neurochemistry 166, 109–137. 10.1111/jnc.15811.

13. Bellesi, M., de Vivo, L., Chini, M., Gilli, F., Tononi, G., and Cirelli, C. (2017). Sleep Loss Promotes Astrocytic Phagocytosis and Microglial Activation in Mouse Cerebral Cortex. J Neurosci 37, 5263–5273. 10.1523/jneurosci.3981-16.2017.

14. Florian, C., Vecsey, C.G., Halassa, M.M., Haydon, P.G., and Abel, T. (2011). Astrocyte-derived adenosine and A1 receptor activity contribute to sleep loss-induced deficits in hippocampal synaptic plasticity and memory in mice. J Neurosci 31, 6956–6962. 10.1523/JNEUROSCI.5761-10.2011.

15. Petit, J.M., and Magistretti, P.J. (2016). Regulation of neuron–astrocyte metabolic coupling across the sleep–wake cycle. Neuroscience 323, 135–156. 10.1016/j.neuroscience.2015.12.007.

16. Haynes, P.R., Pyfrom, E.S., Li, Y., Stein, C., Cuddapah, V.A., Jacobs, J.A., Yue, Z., and Sehgal, A. (2024). A neuron-glia lipid metabolic cycle couples daily sleep to mitochondrial homeostasis. Nat Neurosci 27, 666–678. https://www.nature.com/articles/s41593-023-01568-1.

17. Ingiosi, A.M., Hayworth, C.R., Harvey, D.O., Singletary, K.G., Rempe, M.J., Wisor, J.P., and Frank, M.G. (2020). A Role for Astroglial Calcium in Mammalian Sleep and Sleep Regulation. Current Biology 30, 4373–4383.e4377. 10.1016/j.cub.2020.08.052.

18. Ingiosi, A.M., Hayworth, C.R., and Frank, M.G. (2023). Activation of Basal Forebrain Astrocytes Induces Wakefulness without Compensatory Changes in Sleep Drive. The Journal of Neuroscience 43, 5792–5809. 10.1523/jneurosci.0163-23.2023.

19. Almeida, R.G., and Lyons, D.A. (2014). On the resemblance of synapse formation and CNS myelination. Neuroscience 276, 98–108. 10.1016/j.neuroscience.2013.08.062.

20. de Vivo, L., and Bellesi, M. (2019). The role of sleep and wakefulness in myelin plasticity. Glia 67, 2142–2152. 10.1002/glia.23667.

21. Molina-Gonzalez, I., Holloway, R.K., Jiwaji, Z., Dando, O., Kent, S.A., Emelianova, K., Lloyd, A.F., Forbes, L.H., Mahmood, A., Skripuletz, T., et al. (2023). Astrocyte-oligodendrocyte interaction regulates central nervous system regeneration. Nat Commun 14, 3372. 10.1038/s41467-023-39046-8.

22. Cirelli, C., Gutierrez, C.M., and Tononi, G. (2004). Extensive and divergent effects of sleep and wakefulness on brain gene expression. Neuron 41, 35–43. 10.1016/s0896-6273(03)00814-6.

23. Graves, L.A., Heller, E.A., Pack, A.I., and Abel, T. (2003). Sleep deprivation selectively impairs memory consolidation for contextual fear conditioning. Learn Mem 10, 168–176. 10.1101/lm.48803.

24. Ognjanovski, N., Maruyama, D., Lashner, N., Zochowski, M., and Aton, S.J. (2014). CA1 hippocampal network activity changes during sleep-dependent memory consolidation. Front Syst Neurosci 8, 61. 10.3389/fnsys.2014.00061.

25. Vecsey, C.G., Baillie, G.S., Jaganath, D., Havekes, R., Daniels, A., Wimmer, M., Huang, T., Brown, K.M., Li, X.Y., Descalzi, G., et al. (2009). Sleep deprivation impairs cAMP signalling in the hippocampus. Nature 461, 1122–1125. 10.1038/nature08488.

26. Giri, B., Kinsky, N., Kaya, U., Maboudi, K., Abel, T., and Diba, K. (2024). Sleep loss diminishes hippocampal reactivation and replay. Nature 10.1038/s41586-024-07538-2.

27. Muzur, A., Pace-Schott, E.F., and Hobson, J.A. (2002). The prefrontal cortex in sleep. Trends in Cognitive Sciences 6, 475–481. 10.1016/S1364-6613(02)01992-7.

28. Sigurdsson, T., and Duvarci, S. (2015). Hippocampal-Prefrontal Interactions in Cognition, Behavior and Psychiatric Disease. Front Syst Neurosci 9, 190. 10.3389/fnsys.2015.00190.

29. Mo, A., Mukamel, Eran A., Davis, Fred P., Luo, C., Henry, Gilbert L., Picard, S., Urich, Mark A., Nery, Joseph R., Sejnowski, Terrence J., Lister, R., et al. (2015). Epigenomic Signatures of Neuronal Diversity in the Mammalian Brain. Neuron 86, 1369–1384. 10.1016/j.neuron.2015.05.018.

30. Araque, A., Carmignoto, G., Haydon, P.G., Oliet, S.H., Robitaille, R., and Volterra, A. (2014). Gliotransmitters travel in time and space. Neuron 81, 728–739. 10.1016/j.neuron.2014.02.007.

31. Goenaga, J., Araque, A., Kofuji, P., and Herrera Moro Chao, D. (2023). Calcium signaling in astrocytes and gliotransmitter release. Frontiers in Synaptic Neuroscience Volume 15 - 2023. 10.3389/fnsyn.2023.1138577.

32. Schubert, V., Bouvier, D., and Volterra, A. (2011). SNARE protein expression in synaptic terminals and astrocytes in the adult hippocampus: A comparative analysis. Glia 59, 1472–1488. 10.1002/glia.21190.

33. Jellinger, K.A. (2010). Basic mechanisms of neurodegeneration: a critical update. J Cell Mol Med 14, 457–487. 10.1111/j.1582-4934.2010.01010.x.

34. Vanrobaeys, Y., Peterson, Z.J., Walsh, E.N., Chatterjee, S., Lin, L.-C., Lyons, L.C., Nickl-Jockschat, T., and Abel, T. (2023). Spatial transcriptomics reveals unique gene expression changes in different brain regions after sleep deprivation. Nature Communications 14, 7095. 10.1038/s41467-023-42751-z.

35. Chen, H.Y., Kelley, R.A., Li, T., and Swaroop, A. (2021). Primary cilia biogenesis and associated retinal ciliopathies. Seminars in Cell & Developmental Biology 110, 70–88. 10.1016/j.semcdb.2020.07.013.

36. Liu, S., Trupiano, M.X., Simon, J., Guo, J., and Anton, E.S. (2021). The essential role of primary cilia in cerebral cortical development and disorders. Curr Top Dev Biol 142, 99–146. 10.1016/bs.ctdb.2020.11.003.

37. Tu, H.-Q., Li, S., Xu, Y.-L., Zhang, Y.-C., Li, P.-Y., Liang, L.-Y., Song, G.-P., Jian, X.-X., Wu, M., Song, Z.-Q., et al. (2023). Rhythmic cilia changes support SCN neuron coherence in circadian clock. Science 380, 972–979. doi:10.1126/science.abm1962.

38. Nakazato, R., Matsuda, Y., Ijaz, F., and Ikegami, K. (2023). Circadian oscillation in primary cilium length by clock genes regulates fibroblast cell migration. EMBO reports 24, e56870. 10.15252/embr.202356870.

39. Cardona-Quiñones, R.A., Ramírez-Rivera, E., Álvarez-Torres, E., Salem-Hernández, S.A., Vargas-Pérez, N.J., and De Jesús-Rojas, W. (2025). A Pilot Study of Primary Ciliary Dyskinesia: Sleep-Related Disorders and Neuropsychiatric Comorbidities. Journal of Clinical Medicine 14, 1353.

40. Strobel, M.R., Zhou, Y., Qiu, L., Hofer, A.M., and Chen, X. (2025). Temporal ablation of the ciliary protein IFT88 alters normal brainwave patterns. Scientific Reports 15, 347. 10.1038/s41598-024-83432-1.

41. Oktem, S., Karadag, B., Erdem, E., Gokdemir, Y., Karakoc, F., Dagli, E., and Ersu, R. (2013). Sleep disordered breathing in patients with primary ciliary dyskinesia. Pediatric Pulmonology 48, 897–903. 10.1002/ppul.22710.

42. Han, X., Wei, Y., Wang, H., Wang, F., Ju, Z., and Li, T. (2018). Nonsense-mediated mRNA decay: a ’nonsense’ pathway makes sense in stem cell biology. Nucleic Acids Res 46, 1038–1051. 10.1093/nar/gkx1272.

43. Kurosaki, T., Miyoshi, K., Myers, J.R., and Maquat, L.E. (2018). NMD-degradome sequencing reveals ribosome-bound intermediates with 3 ′ -end non-templated nucleotides. Nature Structural & Molecular Biology 25, 940–950. 10.1038/s41594-018-0132-7.

44. Gaine, M.E., Bahl, E., Chatterjee, S., Michaelson, J.J., Abel, T., and Lyons, L.C. (2021). Altered hippocampal transcriptome dynamics following sleep deprivation. Molecular Brain 14, 125. 10.1186/s13041-021-00835-1.

45. Ma, X., Lu, C., Chen, Y., Li, S., Ma, N., Tao, X., Li, Y., Wang, J., Zhou, M., Yan, Y.-B., et al. (2022). CCT2 is an aggrephagy receptor for clearance of solid protein aggregates. Cell 185, 1325–1345.e1322. 10.1016/j.cell.2022.03.005.

46. Li, J., and Monk, K.R. (2019). Healthy attachments: Cell adhesion molecules collectively control myelin integrity. J Cell Biol 218, 2824–2825. 10.1083/jcb.201907077.

47. Hatta, K., Okada, T.S., and Takeichi, M. (1985). A monoclonal antibody disrupting calcium-dependent cell-cell adhesion of brain tissues: possible role of its target antigen in animal pattern formation. Proceedings of the National Academy of Sciences 82, 2789–2793. doi:10.1073/pnas.82.9.2789.

48. Schnädelbach, O., Ozen, I., Blaschuk, O.W., Meyer, R.L., and Fawcett, J.W. (2001). N-cadherin is involved in axon-oligodendrocyte contact and myelination. Mol Cell Neurosci 17, 1084–1093. 10.1006/mcne.2001.0961.

49. Nakashiba, T., Young, J.Z., McHugh, T.J., Buhl, D.L., and Tonegawa, S. (2008). Transgenic inhibition of synaptic transmission reveals role of CA3 output in hippocampal learning. Science 319, 1260–1264. 10.1126/science.1151120.

50. Guo, J., Oliveros, H.C., Oh, S.J., Liang, B., Li, Y., Kavalali, E.T., Lin, D.-T., and Xu, W. (2021). Stratum Lacunosum-moleculare Interneurons of the Hippocampus Coordinate Memory Encoding and Retrieval. bioRxiv, 2021.2006.2011.448103. 10.1101/2021.06.11.448103.

51. Munyeshyaka, M., and Fields, R.D. (2022). Oligodendroglia are emerging players in several forms of learning and memory. Commun Biol 5, 1148. 10.1038/s42003-022-04116-y.

52. Ingiosi, A.M., and Frank, M.G. (2023). Goodnight, astrocyte: waking up to astroglial mechanisms in sleep. The FEBS Journal 290, 2553–2564. 10.1111/febs.16424.

53. Carvalhas-Almeida, C., and Sehgal, A. (2025). Glia: the cellular glue that binds circadian rhythms and sleep. Sleep 48 10.1093/sleep/zsae314.

54. Aho, V., Ollila, H.M., Kronholm, E., Bondia-Pons, I., Soininen, P., Kangas, A.J., Hilvo, M., Seppälä, I., Kettunen, J., Oikonen, M., et al. (2016). Prolonged sleep restriction induces changes in pathways involved in cholesterol metabolism and inflammatory responses. Scientific Reports 6, 24828. 10.1038/srep24828.

55. Ford, K., Zuin, E., Righelli, D., Medina, E., Schoch, H., Singletary, K., Muheim, C., Frank, M.G., Hicks, S.C., Risso, D., and Peixoto, L. (2024). A global transcriptional atlas of the effect of acute sleep deprivation in the mouse frontal cortex. iScience 27, 110752. 10.1016/j.isci.2024.110752.

56. Jha, P.K., Valekunja, U.K., Ray, S., Nollet, M., and Reddy, A.B. (2022). Single-cell transcriptomics and cell-specific proteomics reveals molecular signatures of sleep. Communications Biology 5, 846. 10.1038/s42003-022-03800-3.

57. Kim, S.J., Hotta-Hirashima, N., Asano, F., Kitazono, T., Iwasaki, K., Nakata, S., Komiya, H., Asama, N., Matsuoka, T., Fujiyama, T., et al. (2022). Kinase signalling in excitatory neurons regulates sleep quantity and depth. Nature 612, 512–518. 10.1038/s41586-022-05450-1.

58. Nakata, S., Iwasaki, K., Funato, H., Yanagisawa, M., and Ozaki, H. (2024). Neuronal subtype-specific transcriptomic changes in the cerebral neocortex associated with sleep pressure. Neuroscience Research. 10.1016/j.neures.2024.03.004.

59. Guo, X., Keenan, B.T., Reiner, B.C., Lian, J., and Pack, A.I. (2024). Single-nucleus RNA-seq identifies one galanin neuronal subtype in mouse preoptic hypothalamus activated during recovery from sleep deprivation. Cell Rep 43, 114192. 10.1016/j.celrep.2024.114192.

60. Beard, E., Lengacher, S., Dias, S., Magistretti, P.J., and Finsterwald, C. (2022). Astrocytes as Key Regulators of Brain Energy Metabolism: New Therapeutic Perspectives. Frontiers in Physiology Volume 12 - 2021. 10.3389/fphys.2021.825816.

61. Calì, C., Cantando, I., Veloz Castillo, M.F., Gonzalez, L., and Bezzi, P. (2024). Metabolic Reprogramming of Astrocytes in Pathological Conditions: Implications for Neurodegenerative Diseases. International Journal of Molecular Sciences 25, 8922.

62. Ferris, H.A., Perry, R.J., Moreira, G.V., Shulman, G.I., Horton, J.D., and Kahn, C.R. (2017). Loss of astrocyte cholesterol synthesis disrupts neuronal function and alters whole-body metabolism. Proc Natl Acad Sci U S A 114, 1189–1194. 10.1073/pnas.1620506114.

63. Zhang, J., and Liu, Q. (2015). Cholesterol metabolism and homeostasis in the brain. Protein & Cell 6, 254–264. 10.1007/s13238-014-0131-3.

64. Poitelon, Y., Kopec, A.M., and Belin, S. (2020). Myelin Fat Facts: An Overview of Lipids and Fatty Acid Metabolism. Cells 9. 10.3390/cells9040812.

65. Wang, H., and Eckel, R.H. (2014). What are lipoproteins doing in the brain? Trends in Endocrinology & Metabolism 25, 8–14. 10.1016/j.tem.2013.10.003.

66. Monje, M. (2018). Myelin Plasticity and Nervous System Function. Annual Review of Neuroscience 41, 61–76. 10.1146/annurev-neuro-080317-061853.

67. Gimpl, G., Burger, K., and Fahrenholz, F. (1997). Cholesterol as Modulator of Receptor Function. Biochemistry 36, 10959–10974. 10.1021/bi963138w.

68. Shin, K.C., Ali Moussa, H.Y., and Park, Y. (2024). Cholesterol imbalance and neurotransmission defects in neurodegeneration. Experimental & Molecular Medicine 56, 1685–1690. 10.1038/s12276-024-01273-4.

69. Vecsey, C.G., Baillie, G.S., Jaganath, D., Havekes, R., Daniels, A., Wimmer, M., Huang, T., Brown, K.M., Li, X.-Y., Descalzi, G., et al. (2009). Sleep deprivation impairs cAMP signalling in the hippocampus. Nature 461, 1122–1125. 10.1038/nature08488.

70. Vecsey, C.G., Huang, T., and Abel, T. (2018). Sleep deprivation impairs synaptic tagging in mouse hippocampal slices. Neurobiology of Learning and Memory 154, 136–140. 10.1016/j.nlm.2018.03.016.

71. Bindocci, E., Savtchouk, I., Liaudet, N., Becker, D., Carriero, G., and Volterra, A. (2017). Three-dimensional Ca^2+^ imaging advances understanding of astrocyte biology. Science 356, eaai8185. doi:10.1126/science.aai8185.

72. Carney, B.N., Illiano, P., Pohl, T.M., Desu, H.L., Mudalegundi, S., Asencor, A.I., Jwala, S., Ascona, M.C., Singh, P.K., Titus, D.J., et al. (2025). Astroglial TNFR2 signaling regulates hippocampal synaptic function and plasticity in a sex dependent manner. bioRxiv, 2025.2003.2013.643110. 10.1101/2025.03.13.643110.

73. Santello, M., Toni, N., and Volterra, A. (2019). Astrocyte function from information processing to cognition and cognitive impairment. Nature Neuroscience 22, 154–166. 10.1038/s41593-018-0325-8.

74. Mauch, D.H., Nagler, K., Schumacher, S., Goritz, C., Muller, E.C., Otto, A., and Pfrieger, F.W. (2001). CNS synaptogenesis promoted by glia-derived cholesterol. Science 294, 1354–1357. 10.1126/science.294.5545.1354.

75. Pfrieger, F.W. (2003). Role of cholesterol in synapse formation and function. Biochimica et Biophysica Acta (BBA) - Biomembranes 1610, 271–280. 10.1016/S0005-2736(03)00024-5.

76. Pfrieger, F.W., and Barres, B.A. (1997). Synaptic Efficacy Enhanced by Glial Cells in Vitro. Science 277, 1684–1687. doi:10.1126/science.277.5332.1684.

77. Ullian, E.M., Sapperstein, S.K., Christopherson, K.S., and Barres, B.A. (2001). Control of synapse number by glia. Science 291, 657–661. 10.1126/science.291.5504.657.

78. Koudinov, A.R., and Koudinova, N.V. (2001). Essential role for cholesterol in synaptic plasticity and neuronal degeneration. FASEB J 15, 1858–1860. 10.1096/fj.00-0815fje.

79. Chen, M., Xu, Y., Huang, R., Huang, Y., Ge, S., and Hu, B. (2017). N-Cadherin is Involved in Neuronal Activity-Dependent Regulation of Myelinating Capacity of Zebrafish Individual Oligodendrocytes In Vivo. Molecular Neurobiology 54, 6917–6930. 10.1007/s12035-016-0233-4.

80. Anghel, L., Ciubară, A., Nechita, A., Nechita, L., Manole, C., Baroiu, L., Ciubară, A.B., and Muat, C.L. (2023). Sleep Disorders Associated with Neurodegenerative Diseases. Diagnostics 13, 2898.

81. Morton, A.J., and Morton, J. (2023). Sleep and Circadian Rhythm Dysfunction in Animal Models of Huntington’s Disease. Journal of Huntington’s Disease 12, 133–148. 10.3233/jhd-230574.

82. Owen, J.E., Zhu, Y., Fenik, P., Zhan, G., Bell, P., Liu, C., and Veasey, S. (2021). Late-in-life neurodegeneration after chronic sleep loss in young adult mice. Sleep 44. 10.1093/sleep/zsab057.

83. Rowe, R.K., Schulz, P., He, P., Mannino, G.S., Opp, M.R., and Sierks, M.R. (2024). Acute sleep deprivation in mice generates protein pathology consistent with neurodegenerative diseases. Front Neurosci 18, 1436966. 10.3389/fnins.2024.1436966.

84. Zhou, G., Liu, S., Yu, X., Zhao, X., Ma, L., and Shan, P. (2019). High prevalence of sleep disorders and behavioral and psychological symptoms of dementia in late-onset Alzheimer disease: A study in Eastern China. Medicine (Baltimore) 98, e18405. 10.1097/MD.0000000000018405.

85. Benito-León, J., Bermejo-Pareja, F., Vega, S., and Louis, E.D. (2009). Total daily sleep duration and the risk of dementia: a prospective population-based study. European Journal of Neurology 16, 990–997. 10.1111/j.1468-1331.2009.02618.x.

86. Zoccolella, S., Savarese, M., Lamberti, P., Manni, R., Pacchetti, C., and Logroscino, G. (2011). Sleep disorders and the natural history of Parkinson’s disease: The contribution of epidemiological studies. Sleep Medicine Reviews 15, 41–50. 10.1016/j.smrv.2010.02.004.

87. Dunot, J., Ribera, A., Pousinha, P.A., and Marie, H. (2023). Spatiotemporal insights of APP function. Current Opinion in Neurobiology 82, 102754. 10.1016/j.conb.2023.102754.

88. Wang, H., Kulas, J.A., Wang, C., Holtzman, D.M., Ferris, H.A., and Hansen, S.B. (2021). Regulation of beta-amyloid production in neurons by astrocyte-derived cholesterol. Proceedings of the National Academy of Sciences 118, e2102191118. doi:10.1073/pnas.2102191118.

89. Wolozin, B. (2004). Cholesterol and the Biology of Alzheimer’s Disease. Neuron 41, 7–10. 10.1016/S0896-6273(03)00840-7.

90. Varma, V.R., Büşra Lüleci, H., Oommen, A.M., Varma, S., Blackshear, C.T., Griswold, M.E., An, Y., Roberts, J.A., O’Brien, R., Pletnikova, O., et al. (2021). Abnormal brain cholesterol homeostasis in Alzheimer’s disease—a targeted metabolomic and transcriptomic study. npj Aging and Mechanisms of Disease 7, 11. 10.1038/s41514-021-00064-9.

91. Murdock, M.H., and Tsai, L.-H. (2023). Insights into Alzheimer’s disease from single-cell genomic approaches. Nature Neuroscience 26, 181–195. 10.1038/s41593-022-01222-2.

92. Al-Dalahmah, O., Sosunov, A.A., Shaik, A., Ofori, K., Liu, Y., Vonsattel, J.P., Adorjan, I., Menon, V., and Goldman, J.E. (2020). Single-nucleus RNA-seq identifies Huntington disease astrocyte states. Acta Neuropathologica Communications 8, 19. 10.1186/s40478-020-0880-6.

93. Auluck, P.K., Caraveo, G., and Lindquist, S. (2010). α-Synuclein: Membrane Interactions and Toxicity in Parkinson’s Disease. Annual Review of Cell and Developmental Biology 26, 211–233. 10.1146/annurev.cellbio.042308.113313.

94. del Toro, D., Xifro, X., Pol, A., Humbert, S., Saudou, F., Canals, J.M., and Alberch, J. (2010). Altered cholesterol homeostasis contributes to enhanced excitotoxicity in Huntington’s disease. J Neurochem 115, 153–167. 10.1111/j.1471-4159.2010.06912.x.

95. Di Pardo, A., Monyror, J., Morales, L.C., Kadam, V., Lingrell, S., Maglione, V., Wozniak, R.W., and Sipione, S. (2019). Mutant huntingtin interacts with the sterol regulatory element-binding proteins and impairs their nuclear import. Human Molecular Genetics 29, 418–431. 10.1093/hmg/ddz298.

96. Kacher, R., Mounier, C., Caboche, J., and Betuing, S. (2022). Altered Cholesterol Homeostasis in Huntington’s Disease. Front Aging Neurosci 14, 797220. 10.3389/fnagi.2022.797220.

97. Lin, C.-H., Lin, H.-I., Chen, M.-L., Lai, T.-T., Cao, L.-P., Farrer, M.J., Wu, R.-M., and Chien, C.-T. (2016). Lovastatin protects neurite degeneration in LRRK2-G2019S parkinsonism through activating the Akt/Nrf pathway and inhibiting GSK3β activity. Human Molecular Genetics 25, 1965–1978. 10.1093/hmg/ddw068.

98. Sipione, S., Rigamonti, D., Valenza, M., Zuccato, C., Conti, L., Pritchard, J., Kooperberg, C., Olson, J.M., and Cattaneo, E. (2002). Early transcriptional profiles in huntingtin-inducible striatal cells by microarray analyses. Human Molecular Genetics 11, 1953–1965. 10.1093/hmg/11.17.1953.

99. Valenza, M., Rigamonti, D., Goffredo, D., Zuccato, C., Fenu, S., Jamot, L., Strand, A., Tarditi, A., Woodman, B., Racchi, M., et al. (2005). Dysfunction of the Cholesterol Biosynthetic Pathway in Huntington’s Disease. The Journal of Neuroscience 25, 9932–9939. 10.1523/jneurosci.3355-05.2005.

100. Galvagnion, C., Buell, A.K., Meisl, G., Michaels, T.C.T., Vendruscolo, M., Knowles, T.P.J., and Dobson, C.M. (2015). Lipid vesicles trigger α-synuclein aggregation by stimulating primary nucleation. Nature Chemical Biology 11, 229–234. 10.1038/nchembio.1750.

101. Mahul-Mellier, A.-L., Burtscher, J., Maharjan, N., Weerens, L., Croisier, M., Kuttler, F., Leleu, M., Knott, G.W., and Lashuel, H.A. (2020). The process of Lewy body formation, rather than simply α-synuclein fibrillization, is one of the major drivers of neurodegeneration. Proceedings of the National Academy of Sciences 117, 4971–4982. doi:10.1073/pnas.1913904117.

102. Verweij, I.M., Romeijn, N., Smit, D.J.A., Piantoni, G., Van Someren, E.J.W., and van der Werf, Y.D. (2014). Sleep deprivation leads to a loss of functional connectivity in frontal brain regions. BMC Neuroscience 15, 88. 10.1186/1471-2202-15-88.

103. Weng, X., Wen, K., Guo, J., Zhang, P., Zhang, Y., Cao, Q., Han, Q., and Xu, F. (2025). The impact of sleep deprivation on the functional connectivity of visual-related brain regions. Sleep Medicine 125, 155–167. 10.1016/j.sleep.2024.11.026.

104. Wang, X., and Michaelis, E.K. (2010). Selective neuronal vulnerability to oxidative stress in the brain. Frontiers in Aging Neuroscience Volume 2 - 2010. 10.3389/fnagi.2010.00012.

105. Wang, L., Park, L., Wu, W., King, D., Vega-Medina, A., Raven, F., Martinez, J., Ensing, A., McDonald, K., Yang, Z., et al. (2024). Sleep-dependent engram reactivation during hippocampal memory consolidation associated with subregion-specific biosynthetic changes. iScience 27, 109408. 10.1016/j.isci.2024.109408.

106. Goicoechea, L., Conde de la Rosa, L., Torres, S., García-Ruiz, C., and Fernández-Checa, J.C. (2023). Mitochondrial cholesterol: Metabolism and impact on redox biology and disease. Redox Biology 61, 102643. 10.1016/j.redox.2023.102643.

107. Chiu, Y.-C., Chu, P.-W., Lin, H.-C., and Chen, S.-K. (2021). Accumulation of cholesterol suppresses oxidative phosphorylation and altered responses to inflammatory stimuli of macrophages. Biochemistry and Biophysics Reports 28, 101166. 10.1016/j.bbrep.2021.101166.

108. Solsona-Vilarrasa, E., Fucho, R., Torres, S., Nuñez, S., Nuño-Lámbarri, N., Enrich, C., García-Ruiz, C., and Fernández-Checa, J.C. (2019). Cholesterol enrichment in liver mitochondria impairs oxidative phosphorylation and disrupts the assembly of respiratory supercomplexes. Redox Biology 24, 101214. 10.1016/j.redox.2019.101214.

109. Sarnataro, R., Velasco, C.D., Monaco, N., Kempf, A., and Miesenböck, G. (2025). Mitochondrial origins of the pressure to sleep. Nature 645, 722–728. 10.1038/s41586-025-09261-y.

110. Sterpka, A., and Chen, X. (2018). Neuronal and astrocytic primary cilia in the mature brain. Pharmacological Research 137, 114–121. 10.1016/j.phrs.2018.10.002.

111. Wheway, G., Nazlamova, L., and Hancock, J.T. (2018). Signaling through the Primary Cilium. Frontiers in Cell and Developmental Biology Volume 6 - 2018. 10.3389/fcell.2018.00008.

112. Mill, P., Christensen, S.T., and Pedersen, L.B. (2023). Primary cilia as dynamic and diverse signalling hubs in development and disease. Nature Reviews Genetics 24, 421–441. 10.1038/s41576-023-00587-9.

113. Hasenpusch-Theil, K., and Theil, T. (2021). The Multifaceted Roles of Primary Cilia in the Development of the Cerebral Cortex. Frontiers in Cell and Developmental Biology Volume 9 - 2021. 10.3389/fcell.2021.630161.

114. Alvarez Retuerto, A.I., Cantor, R.M., Gleeson, J.G., Ustaszewska, A., Schackwitz, W.S., Pennacchio, L.A., and Geschwind, D.H. (2008). Association of common variants in the Joubert syndrome gene (AHI1) with autism. Hum Mol Genet 17, 3887–3896. 10.1093/hmg/ddn291.

115. Migliavacca, E., Golzio, C., Mannik, K., Blumenthal, I., Oh, E.C., Harewood, L., Kosmicki, J.A., Loviglio, M.N., Giannuzzi, G., Hippolyte, L., et al. (2015). A Potential Contributory Role for Ciliary Dysfunction in the 16p11.2 600 kb BP4-BP5 Pathology. Am J Hum Genet 96, 784–796. 10.1016/j.ajhg.2015.04.002.

116. Guemez-Gamboa, A., Coufal, Nicole G., and Gleeson, Joseph G. (2014). Primary Cilia in the Developing and Mature Brain. Neuron 82, 511–521. 10.1016/j.neuron.2014.04.024.

117. Valente, E.M., Rosti, R.O., Gibbs, E., and Gleeson, J.G. (2014). Primary cilia in neurodevelopmental disorders. Nature Reviews Neurology 10, 27–36. 10.1038/nrneurol.2013.247.

118. Alhassen, W., Chen, S., Vawter, M., Robbins, B.K., Nguyen, H., Myint, T.N., Saito, Y., Schulmann, A., Nauli, S.M., Civelli, O., et al. (2021). Patterns of cilia gene dysregulations in major psychiatric disorders. Progress in Neuro-Psychopharmacology and Biological Psychiatry 109, 110255. 10.1016/j.pnpbp.2021.110255.

119. Muhamad, N.A., Masutani, K., Furukawa, S., Yuri, S., Toriyama, M., Matsumoto, C., Itoh, S., Shinagawa, Y., Isotani, A., Toriyama, M., and Itoh, H. (2024). Astrocyte-Specific Inhibition of the Primary Cilium Suppresses C3 Expression in Reactive Astrocyte. Cellular and Molecular Neurobiology 44, 48. 10.1007/s10571-024-01482-5.

120. Schartz, N.D., and Tenner, A.J. (2020). The good, the bad, and the opportunities of the complement system in neurodegenerative disease. Journal of Neuroinflammation 17, 354. 10.1186/s12974-020-02024-8.

121. Miyamoto, T., Hosoba, K., Itabashi, T., Iwane, A.H., Akutsu, S.N., Ochiai, H., Saito, Y., Yamamoto, T., and Matsuura, S. (2020). Insufficiency of ciliary cholesterol in hereditary Zellweger syndrome. The EMBO Journal 39, e103499. 10.15252/embj.2019103499.

122. Walsh, E.N., Shetty, M.S., Diba, K., and Abel, T. (2023). Chemogenetic Enhancement of cAMP Signaling Renders Hippocampal Synaptic Plasticity Resilient to the Impact of Acute Sleep Deprivation. eNeuro 10. 10.1523/ENEURO.0380-22.2022.

